# An expanded evolutionary landscape of H3 histones with variants that destabilize nucleosomes and promote chromatin accessibility

**DOI:** 10.64898/2026.11.04.764309

**Authors:** Carlos F. Rivera-López, Michael L. Nosella, Siddhartha G. Jena, Paul Bump, Elizabeth M. Duncan, Jason D. Buenrostro, Karolin Luger, Mansi Srivastava

## Abstract

Histones are among the most conserved proteins in eukaryotes, yet many animal lineages have evolved divergent histone variants with specialized functions. Some broadly conserved variants (e.g., H3.3) have been linked to the regulation of chromatin accessibility in stem cells, but whether highly divergent variants function similarly remains unclear. Here, we characterize the histone repertoire of the acoel *Hofstenia miamia*, an animal that can undergo whole-body regeneration using adult pluripotent stem cells (aPSCs). We identify two novel histone H3 paralogs, H3-a and H3-b, that are expressed in aPSCs and form a lineage-specific sister clade to H3.2 and H3.3. Despite substantial sequence divergence and reduced compatibility with canonical chaperone machinery, both variants assemble bona fide nucleosomes *in vitro*. These nucleosomes exhibit reduced stability and incomplete DNA wrapping, and a cryo-EM structure of the H3-b nucleosome reveals the disruption of conserved histone-DNA contacts. *In vivo*, H3-a and H3-b are upregulated following injury, and their depletion reduces genome-wide chromatin accessibility, suggesting a role in the maintenance of accessible chromatin during regeneration. A survey of H3 sequences across animal phyla shows that H3-a and H3-b belong to a broader landscape of highly divergent H3 proteins, some of which exhibit sequence differences across both the N-terminal tail and histone-fold domain. Together, these findings link histone sequence diversification to chromatin regulation in aPSCs and highlight the underappreciated evolutionary diversity and functional potential of divergent animal H3 proteins.

## INTRODUCTION

Histone proteins are essential components of chromatin that assemble into nucleosomes to package eukaryotic DNA and regulate genome accessibility^1^ (Fig. 1A). Histones can be categorized as ‘canonical’, with expression during S phase to package newly synthesized DNA, or ‘variant’, with replication-independent expression and specialized roles in transcription, DNA repair, and chromatin architecture^2^. Both canonical and variant histones are highly conserved across eukaryotic evolution, but recent work leveraging expanded genomic datasets has revealed that metazoans have explored a broader range of histone sequences beyond the well-conserved repertoire^3–12^. Although some divergent histones have been implicated in specialized functions, including DNA packaging in sperm and maintenance of genome integrity in cnidarians^12,13^, the extent of histone sequence diversification across metazoans and their functional significance remain poorly characterized. Here, we combine *in vitro* structural and biochemical approaches with *in vivo* functional studies to characterize two novel histone H3 proteins, providing experimental insight into a large but previously unstudied class of animal histones.

**Figure 1.**
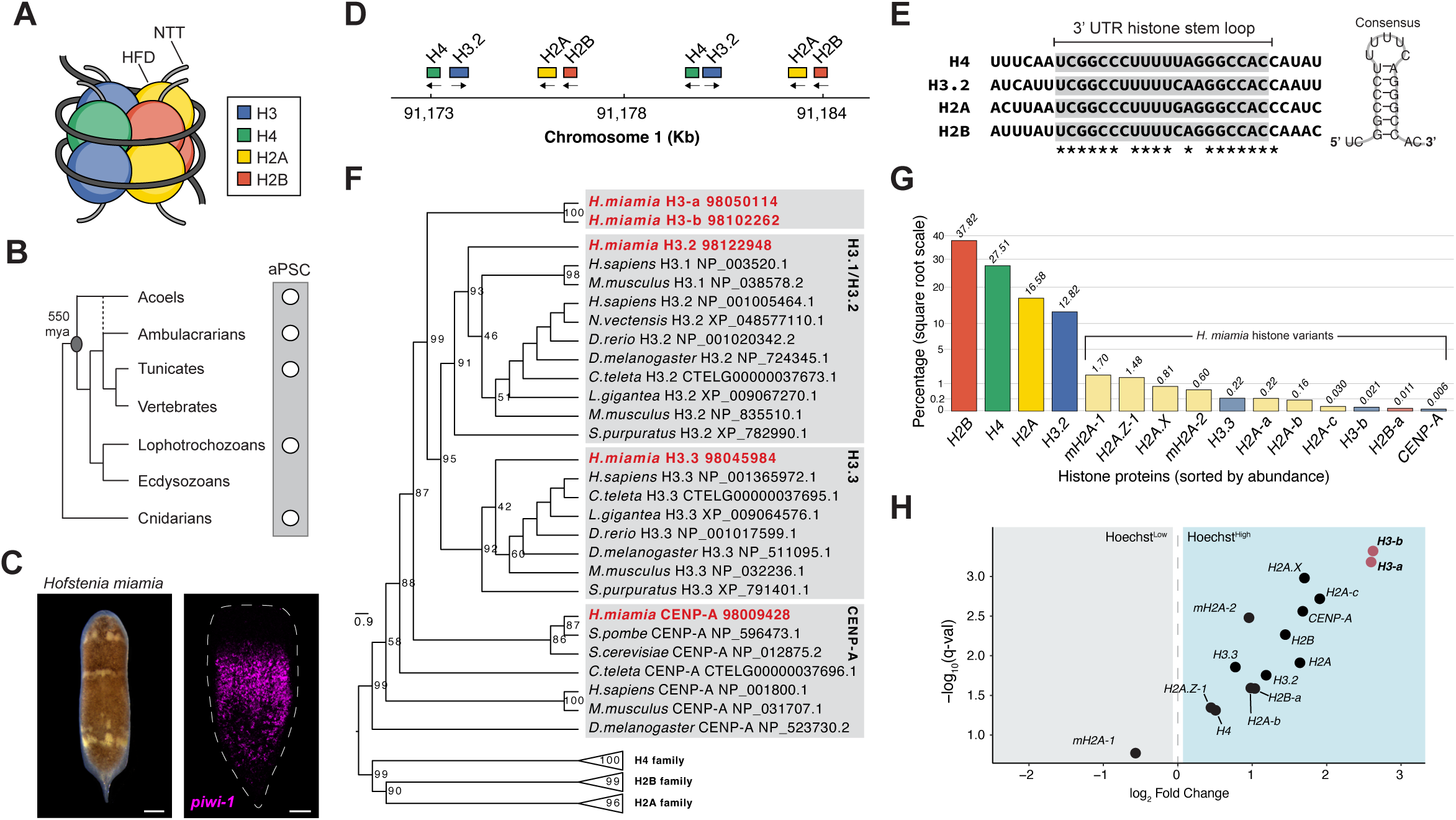
*Hofstenia miamia* encodes two novel histone H3 proteins. (A) Schematic of a canonical nucleosome, in which histone fold domains (HFDs) assemble into the histone octamer around which DNA is wrapped in two gyres. N-terminal tails (NTTs) extend outward from the nucleosome core. Histone proteins are color-coded: H3, blue; H4, green; H2A, yellow; H2B, red. (B) Schematic phylogeny of bilaterian animal groups highlighting the presence of aPSCs across multiple metazoan lineages and the origin of bilateria 550 million years ago (mya). The dashed line represents alternate placement of Xenacoelomorpha^116^. (C) *H. miamia* juvenile and a fluorescent *in situ* hybridization (FISH) representative image of *piwi-1*, marker of adult pluripotent stem cells (aPSC). Scale bars, 250 and 100 µm, respectively. (D) Examples of clustered units of canonical histone genes in chromosome 1. Directionality of each gene is represented by arrows. (E) Multiple sequence alignment of histone 3′ UTR stem-loop regions with conserved positions indicated by asterisks (left), and a schematic of the derived consensus secondary structure (right). (F) Maximum-likelihood phylogenetic tree of histone H3 homologs with bootstrap support values shown at each node. *H. miamia* sequences are highlighted in red: *H3.2*, *H3.3* and *CENP-A* cluster with their respective homologs, whereas *H3-a* and *H3-b* form a distinct sister clade to the *H3.2* and *H3.3* groups. H2A, H2B, and H4 families are shown as collapsed clades. (G) Mass spectrometry-based quantification of histone proteins purified from *H. miamia* nuclei. Square root scale percentages of detected canonical and variant histones. (H) Volcano plot of bulk differential expression between populations with low or high Hoechst staining, where each point represents a histone gene^44^. *H3-a* and *H3-b* are highlighted in red and display the strongest enrichment within the Hoechst^high^ cells.

Acoel worms diverged from other bilaterians ∼550 million years ago and represent a phylum (Xenacoelomorpha) that informs the evolution of metazoan molecular and cellular processes (Fig. 1B). The marine worm *Hofstenia miamia* is a new acoel model organism that is amenable to mechanistic studies of regeneration. Whole-body regeneration in *H. miamia* relies on a population of adult pluripotent stem cells (aPSCs), also known as neoblasts, that express *piwi-1* and other germline multipotency program genes^14^ (Fig. 1C). Similar *piwi*-expressing cells have been described across metazoans, including in sponges, cnidarians, planarians, annelids, and tunicates^15,16^ (Fig. 1B), raising the possibility that aPSCs represent an ancient animal cell type. *H. miamia* aPSCs are transcriptionally heterogeneous, consisting of many lineage-primed subpopulations that are hypothesized to give rise to their associated tissues^17^. One subpopulation, however, lacks markers of lineage commitment and exhibits high *piwi-1* expression, consistent with a more unspecialized aPSC state. Also enriched within this subpopulation are two highly diverged H3 variants, presenting an opportunity to study novel H3 variants with potential roles in regulating stem cell biology.

aPSCs likely require a chromatin state that enables access to a broad repertoire of gene programs as they differentiate into diverse lineages. Canonical histones and their conserved variants have been implicated in regulating stem cell identity and differentiation in several species. In planarians, which also undergo aPSC-based whole-body regeneration, canonical histones have been linked to neoblast maintenance and differentiation^18,19^, but the roles of histone variants have remained unexplored. During vertebrate embryonic development, the histone variants H2A.Z and H3.3 maintain pluripotency of cells by tuning chromatin accessibility at promoters and enhancers of pluripotency-associated genes^20–24^. As these cells differentiate, H3.3 is redistributed from pluripotency-associated genes to lineage-specific loci that drive maturation^25–29^. These and many other studies in vertebrates focus heavily on understanding highly conserved histone variants, leaving open the question of how more divergent histones could regulate chromatin in stem cells.

Here, we systematically identified and annotated core and variant histone genes in the *Hofstenia miamia* genome and confirmed their presence at the protein level. Focusing on members of the histone H3 family, we characterized two highly divergent variants: H3-a and H3-b. We determined that these divergent histone proteins assemble into nucleosomes with reduced stability and incomplete DNA wrapping, associated with weakened interactions between the histone octamer and nucleosomal DNA. *In vivo* experiments investigating the role of these variants in homeostatic tissue turnover and regeneration revealed a functional role for *H3-a* and *H3-b* in promoting chromatin accessibility. Extending our analysis across animal phyla, we found that H3-a and H3-b belong to a previously functionally uncharacterized class of highly divergent H3 proteins distributed across animal lineages. Thus, by leveraging a highly regenerative research organism, this work links histone sequence diversification to chromatin regulation in aPSCs and provides a framework for understanding how histone variants may contribute to chromatin states across metazoans.

## RESULTS

### The *Hofstenia miamia* genome encodes conserved canonical and variant histone proteins, as well as novel histone variants

Building on the previous identification of two H3 histone variants^17^, we characterized the complete repertoire of nucleosomal histone protein diversity in *Hofstenia miamia*. The nucleosome core, an octamer comprising two H2A-H2B dimers and one H3-H4 tetramer, wraps ∼147 base pairs of DNA (Fig. 1A). We used human nucleosomal histones from the NCBI Histone Database 2.0^30^ to identify unique genes encoding members of the H2A, H2B, H3, and H4 histone families in the *H. miamia* genome^31^ (Supplementary File 1). Using a maximum-likelihood phylogenetic reconstruction of known histone proteins from slow-evolving animals, we assigned orthology to each of the identified histone proteins (Supplementary Fig. 1A). Together, this analysis identified *H. miamia* proteins corresponding to both canonical and variant histones across all nucleosomal histone families, indicating conservation of the full histone repertoire^2,32^.

Genes encoding canonical histone proteins (namely, H2A, H2B, H3.2 and H4) display two key features across well-studied organisms: 1) they are present in multiple copies and are organized into several clusters across the genome, and 2) their transcripts contain conserved secondary RNA structures^33,34^. In the *H. miamia* genome, homologs of canonical histone genes are clustered in chromosomes 1, 2, and 8 (Fig. 1D, Supplementary Fig. 1B, and Supplementary File 1). Canonical histone transcripts are subject to post-transcriptional regulation mediated by binding of the stem-loop binding protein (SLBP) to a stem-loop structure that forms in their 3’ untranslated region (UTR)^35,36^. Consistent with this, *H. miamia* canonical histone genes encode a sequence in their 3’ UTR predicted to fold into a homologous stem-loop structure^37,38^ (Fig. 1E, Supplementary File 1), suggesting that transcripts of canonical genes can be a substrate of SLBP binding for gene regulation. The *H. miamia* histone variant H2A.X also forms a 3’ UTR stem-loop, similar to loop formation in other systems^39–41^. The remaining identified histone genes in *H. miamia* encode histone variants, which instead are encoded at unique genomic loci, do not appear to be clustered on specific chromosomes, and lack evidence of 3’ UTR stem-loop structures (Supplementary File 1).

The *H. miamia* histone H3 family contains five members (Fig. 1F and Supplementary Fig. 1A). H3.2 and H3.3 cluster within their respective clades in our phylogenetic analysis and represent the well-characterized replication-dependent and -independent H3 proteins, respectively, while a third member clusters with the centromeric H3 variant CENP-A^42,43^. In contrast, *H. miamia* H3-a and H3-b form a distinct sister clade to the H3.2 and H3.3 subfamilies (Fig. 1F). Although H3-a and H3-b share high nucleotide sequence similarity (91.53% identity, Supplementary File 1), individual PacBio long reads spanning each locus and its flanking genomic regions confirmed that they are encoded at distinct genomic loci and are therefore bona fide paralogs^31^ (Supplementary Fig. 1C). Based on these phylogenetic and genomic features, together with RNA structure properties, we renamed these histone genes, which were previously reported as H3.3, H3.3-2, H3.3-3 and H3.3-4 histones^17^ as H3-a, H3-b, H3.3 and CENP-A, respectively. This improved nomenclature reflects the identity of H3-a and H3-b as an *H. miamia*-specific sister clade to H3.2, H3.3, and CENP-A (Fig. 1F).

Next, we used mass spectrometry to profile the histone proteins present in *H. miamia*. We optimized a trichloroacetic acid-based extraction protocol to purify the endogenous nuclear histone proteins from *H. miamia* juveniles (see Methods). This approach yielded samples enriched for proteins of approximately 10-15 kDa, consistent with the expected molecular weight of histone proteins (Supplementary Fig. 1D), allowing subsequent mass spectrometry to quantify their abundance. As expected, we found that the canonical histones were the most abundant histone proteins detected, whereas histone variants were detected at relatively lower abundances (Fig. 1G).

To assess whether the protein abundances of these histones are reflective of their transcriptional expression levels, we next leveraged available transcriptomic datasets for *H. miamia*. We analyzed bulk RNA-sequencing (RNA-seq) data from Hoechst-defined cell populations^44^, alongside single-cell RNA-sequencing (scRNA-seq) data from intact and regenerating *H. miamia* animals^17^. Given that aPSCs are the only proliferative cells in *H. miamia*, the Hoechst^high^ fraction is enriched for aPSCs, whereas the Hoechst^low^ fraction predominantly contains non-aPSCs. Bulk differential expression analysis showed that histone genes are broadly enriched in the aPSC fraction, consistent with the elevated DNA content that characterizes *piwi-1*+ aPSCs (Fig. 1H, Supplementary File 3). Although most histone genes show enrichment in aPSCs, several are also expressed in non-aPSCs (Supplementary Fig. 1E). To examine this, we utilized single-cell RNA-sequencing (scRNA-seq) to assess the distribution of histone transcripts across cell populations. This analysis confirmed that some variants, including *H3.3*, *H2A.X*, and *H2A.Z*, exhibit broad expression profiles that extend beyond the aPSC population (Supplementary Fig. 1F). These patterns are consistent with previous observations that histone variants can be expressed independently of DNA replication and exhibit distinct cell-type-specific expression profiles^45,46^. In contrast, other histone variants, such as *H2A-a*, *H3-a*, *H3-b*, and *CENP-A*, showed lower transcript levels across major cell populations in the hatchling juvenile dataset. Several histone variants like *H2A-b*, *H2A-c*, and *H2B-a* were associated with the germ line, which matures in late-stage adults, and were correspondingly underrepresented in these juvenile samples (Supplementary Fig. 1F-G). These findings corroborated our proteomic data, indicating that differences in histone abundance are associated with variation in expression across life stages and cell types. Collectively, these genomic, proteomic and transcriptomic analyses demonstrate that both canonical and variant histones are conserved and differentially expressed across cell types in *H. miamia*.

Notably, the *H3-a* and *H3-b* variants exhibited the strongest enrichment in the aPSC fraction among all identified histones (Fig. 1H). Given their observed phylogenetic placement within the H3 family (Fig. 1F), we sought to define the molecular features that distinguish H3-a and H3-b from highly conserved H3 proteins that have been well-studied in model organisms.

### Structural divergence of H3-a and H3-b alters conserved histone features and chaperone compatibility

Each core histone contains a histone fold domain (HFD), which mediates interactions that support nucleosome architecture, and an N-terminal tail (NTT) that contains most of the modifiable residues that regulate chromatin dynamics. Among the core histones, H3 contains the greatest number of modifiable residues, making this histone family and its variants important regulators of local chromatin states^47^. Sequence variation within these domains provides a basis for distinguishing canonical and variant histones within a given family. For example, H3.3 members carry a diagnostic AAIG motif at residues 87-90 in the HFD, relative to SAVM in H3.2, along with serine 31 in place of alanine in the NTT^48^. As the *H. miamia* histone H3-a and H3-b are H3 paralogs that fall outside established H3 subfamilies (Fig. 1F), we analyzed the amino acid residues that make up the HFDs and NTTs of these peptides (Supplementary Fig. 2A).

The histone H3 HFD adopts a canonical helix-loop-helix conformation that mediates histone-histone interactions through heterodimerization and contributes to the assembly of the histone octamer within the nucleosome^1^. Despite substantial sequence divergence relative to H3.2 and H3.3, H3-a and H3-b retain broadly similar amino acid charge distributions and recognizable features of the conserved HFD (Fig. 2A). Sequence-based domain annotation using InterProScan identified fold domains in all *H. miamia* H3 proteins^49^, while AlphaFold2 predicted that H3-a and H3-b retain the characteristic HFD architecture (Fig. 2B, Supplementary Fig. 2A).

**Figure 2.**
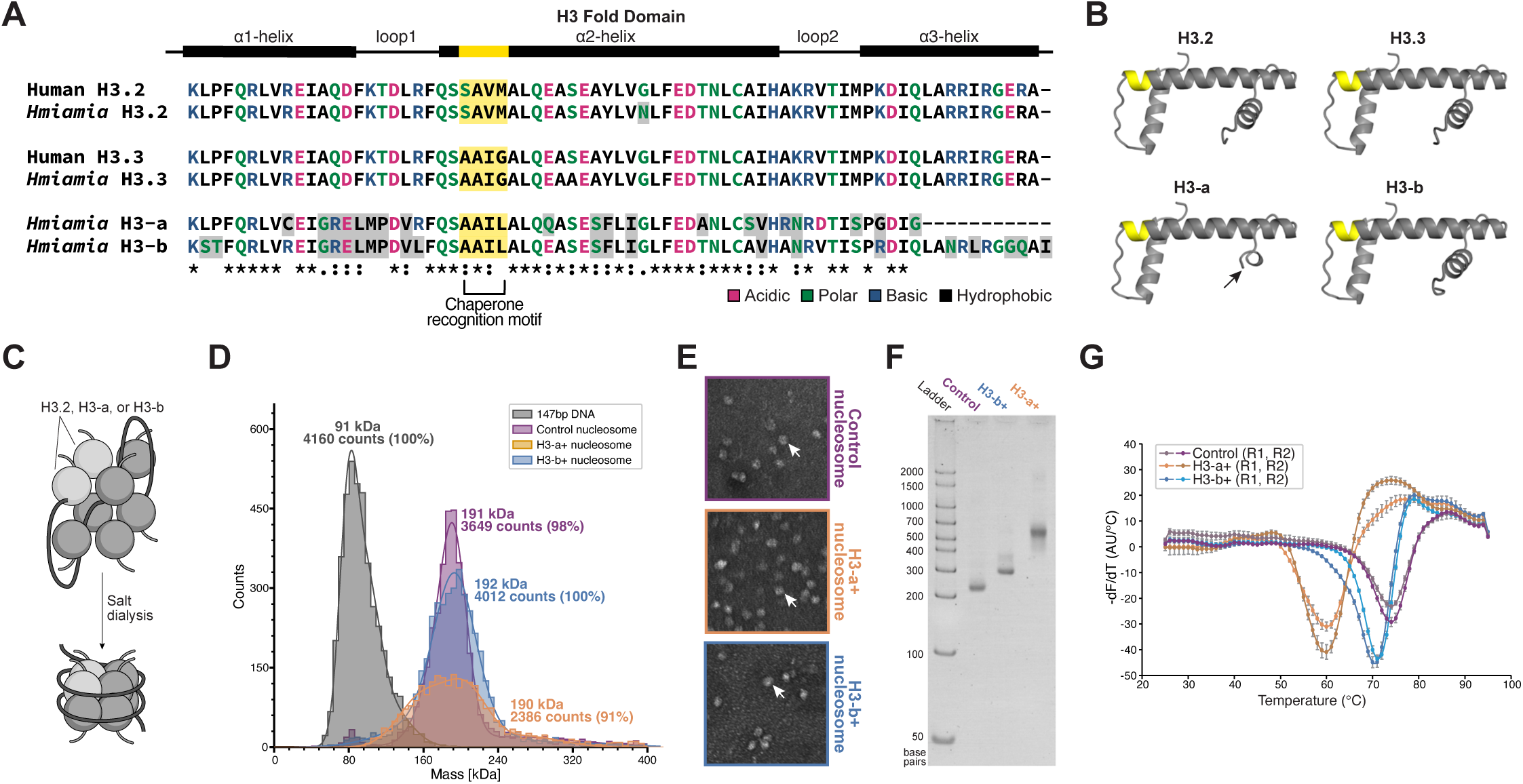
H3-a and H3-b assemble into destabilized nucleosomes *in vitro*. (A) Multiple sequence alignment of the HFD of human and *H. miamia* H3 proteins. ɑ-helices and loop regions are indicated above the alignment. Residues are colored by biochemical properties: acidic (magenta), basic (blue), polar (green), and hydrophobic (black). Residues that diverge from human H3.2 are additionally highlighted with gray shading. The chaperone-recognition motif is highlighted in yellow. (B) AlphaFold2-predicted structures of *H. miamia* H3 HFDs. The chaperone-recognition motif is highlighted in yellow; the black arrow indicates the C-terminal truncation of H3-a. (C) Schematic of *in vitro* nucleosome assembly. H3.2+ (control), H3-a+, or H3-b+ was combined with H2A, H2B, H4 and high-affinity nucleosome-positioning DNA and assembled by salt dialysis. (D) Mass photometry of reconstituted nucleosome complexes. H3-a+ (orange), H3-b+ (blue), and control (purple) nucleosomes formed homogeneous populations with measured molecular masses consistent with the expected nucleosome mass (190-192 kDa). Particle distributions and relative abundances are shown. (E) Negative-stain transmission electron microscopy (TEM) of reconstituted nucleosomes (white arrows). Scale bar, 100 nm. (F) Native polyacrylamide gel electrophoresis analysis of reconstituted nucleosomes. H3-a+ and H3-b+ nucleosomes exhibit reduced electrophoretic mobility relative to control nucleosomes. DNA size markers are indicated in base pairs (bp). (G) Thermal stability of reconstituted nucleosomes measured by fluorescence-based thermal unfolding. Fluorescence changes are plotted as the first derivative (−dF/dT) as a function of temperature.

H3-a and H3-b also harbor a modified chaperone recognition motif (AAIL), distinct from the conserved SAVM or AAIG motifs found in H3.2 and H3.3, respectively, which have been proposed to function as recognition sites for histone-interacting proteins^26,50–53^. To assess the compatibility of H3-a and H3-b with mammalian histone-deposition machinery, we expressed GFP-tagged *H. miamia* histones in a heterologous human NIH-3T3 system^54^. We found that GFP-tagged H3.2 and H3.3 localized strongly to the nucleus and associated with mitotic chromatin in mammalian cells, whereas H3-a and H3-b showed reduced nuclear accumulation and little detectable association with mitotic chromosomes (Supplementary Fig. 2B-E). In contrast, proteomic analyses detected H3-b in *H. miamia* nuclear extracts (Fig. 1G), indicating that this impaired behavior in mammalian cells does not reflect an intrinsic inability to enter the nucleus or to associate with chromatin. Rather, these results suggest that sequence divergence in H3-a and H3-b reduces their compatibility with canonical mammalian histone-deposition machinery. Consistent with the possibility of species-specific deposition mechanisms, several components of replication-independent histone chaperone pathways also show sequence divergence in *H. miamia* (Supplementary Fig. 2F-L, Supplementary File 1).

The NTT of histone H3 proteins contains numerous residues subject to post-translational modifications that contribute to chromatin regulation, while the adjacent ɑN helix participates in histone-DNA interactions that stabilize nucleosomal DNA near its entry and exit sites^47^. Like the majority of histone NTTs, the NTTs of all *H. miamia* H3 proteins were predicted to be intrinsically disordered by MobiDBLite analysis (Supplementary Fig. 2A)^55^. However, H3-a and H3-b exhibit substantial sequence divergence across the N-terminal regions, including substitutions at residues associated with established post-translational modifications. Notably, lysine 4, a conserved site of histone methylation, is replaced by serine in both variants (Supplementary Fig. 2M), eliminating this canonical modification site. Sequence divergence also extends into the adjacent ɑN region, raising the possibility that these variants differ from canonical H3 proteins not only in their potential for post-translational modification but also in their interactions with nucleosomal DNA.

Together, these findings show that H3-a and H3-b retain recognizable H3 structural features while exhibiting sequence properties that distinguish them from canonical H3 proteins and reduce their compatibility with mammalian histone-deposition machinery.

### H3-a and H3-b, together with H2A, H2B, and H4, assemble into nucleosomes *in vitro* with incomplete DNA wrapping

We next asked whether the divergent properties discussed above affect the ability of H3-a and H3-b to assemble into nucleosomes. We performed *in vitro*, chaperone-independent nucleosome reconstitutions by expressing and purifying recombinant H3-a and H3-b proteins from *E. coli* and combining them with human H2A, H2B, and H4, which exhibit high sequence conservation with their *H. miamia* counterparts (Supplementary File 1). As a positive control, we used the human H3.2, which is highly similar to *H. miamia* H3.2 (98% identity, Supplementary File 1). Histone octamers were reconstituted onto a 147-bp DNA fragment containing the Widom 601 nucleosome-positioning sequence, which promotes the formation of homogeneous nucleosomes suitable for downstream biophysical analyses^56,57^ (Fig. 2C). Histone complexes containing H3-a or H3-b formed discrete protein-DNA assemblies and were capable of saturating the 601 DNA substrate (Supplementary Fig. 2N).

We further assessed the composition of the reconstituted assemblies using label-free mass photometry, which determines molecular mass based on light-scattering properties^58,59^. H3-a- and H3-b-containing assemblies exhibited masses consistent with nucleosomes containing a complete histone octamer and 601 DNA, with the predominant populations exhibiting masses consistent with complete nucleosomes and limited evidence of free DNA or subnucleosomal assemblies (Fig. 2D and Supplementary Fig. 2O). Notably, H3-a-containing nucleosomes exhibited a broader mass distribution than H3-b and control nucleosomes, suggesting greater heterogeneity within the reconstituted population. As a negative control, we performed reconstitutions lacking H2A/H2B dimers, which yielded heterogeneous histone-DNA complexes alongside free DNA, consistent with incomplete nucleosome assembly (Supplementary Fig. 2N-P).

To assess the morphology of these assemblies, we visualized reconstituted nucleosomes by negative-stain transmission electron microscopy (TEM). All nucleosomes assembled with canonical H3.2, H3-a, or H3-b exhibited comparable circular morphologies in TEM micrographs (Fig. 2E and Supplementary Fig. 2Q). Together, these results establish that, despite their sequence divergence, H3-a and H3-b can substitute for canonical H3 within fully assembled nucleosomes.

Having determined that H3-a and H3-b can form nucleosomes, we next asked whether incorporation of these variants alters nucleosome architecture or stability. On native polyacrylamide gels, nucleosomes containing H3-a or H3-b exhibited slower migration than those assembled with canonical H3.2, with a more pronounced shift observed for H3-a-containing nucleosomes (Fig. 2F). Because these species possess similar masses (Fig. 2D), we reasoned that their differential electrophoretic migration may be attributed to altered structural conformation. Notably, the HFD of H3-a is shorter (Fig. 2B), lacking part of the predicted ɑ3 helix known to stabilize the four-helix bundle structure between H3 and H3’ that joins together the two H3-H4 dimers^1^. Given that this interface is important for (H3-H4)_2_ tetramer formation and nucleosome stability^60,61^, this truncation raises the possibility that H3-a alters interactions within the histone core or between the histone octamer and DNA.

We next generated low-resolution density maps from the negative-stain TEM micrographs to investigate whether the altered electrophoretic migration reflected differences in nucleosome architecture (Supplementary Fig. 2R). The reconstructed density map of H3-b nucleosomes broadly resembled the canonical H3.2 nucleosome structure, retaining disk-like configuration with two apparent gyres of DNA wrapped around the histone core. In contrast, the H3-a nucleosome density map accommodated only a single well-resolved DNA gyre, consistent with increased conformational flexibility or reduced wrapping of DNA around the histone octamer. Together, these observations suggest that H3-a-containing nucleosomes exhibit less consistently ordered DNA wrapping, whereas any corresponding structural differences in H3-b nucleosomes are less apparent at this resolution.

To determine whether these differences in nucleosome organization were accompanied by altered stability, we measured the thermal stability of H3.2, H3-a and H3-b nucleosomes using a SYPRO Orange assay. H3-a nucleosomes exhibited the lowest thermal stability, with an unfolding transition at 59 °C, followed by H3-b nucleosomes at approximately 71 °C, with canonical H3.2 nucleosomes exhibiting the greatest stability at approximately 75 °C (Fig. 2G). Thus, incorporation of either variant reduces nucleosome stability relative to canonical H3.2, with the more pronounced destabilization observed for H3-a consistent with its altered histone fold and differences in DNA organization *in vitro*.

Together, these results establish that H3-a and H3-b are bona fide nucleosomal histones that confer distinct biophysical properties onto the assemblies they form, including altered DNA organization and reduced nucleosome stability. These properties are consistent with the potential for H3-a- and H3-b-containing nucleosomes to create a more dynamic chromatin environment by weakening histone-DNA interactions. Given these behaviors *in vitro*, we next turned to *H. miamia* to assess the potential roles of H3-a and H3-b *in vivo* during homeostatic tissue turnover and regeneration.

### H3-a and H3-b exhibit distinct homeostatic expression and a shared regenerative response in aPSCs

In *H. miamia*, *piwi-1*+ aPSCs represent the population of actively cycling cells responsible for both homeostatic tissue turnover and whole-body regeneration^62^. Given the known roles of histone variants in regulating pluripotency during vertebrate embryonic development^20–24^, we studied the expression patterns of *H3-a* and *H3-b* during homeostasis and regeneration.

Within subsetted *piwi-1*+ aPSCs spanning all regenerating timepoints, *H3-a* and *H3-b* were enriched in two subpopulations: one hypothesized to represent the unspecialized aPSC state and another population inferred to represent putative germline-specialized neoblasts based on their enriched expression of the *boule-like* (*boll*) gene homolog^17^ (Fig. 3A-B). Using a different probe from that used in Hulett et al. (2023) (Supplementary Fig. 3A-D, Supplementary File 1-2), we identified two spatially distinct subpopulations of *H3-a*-expressing cells in intact animals: a subset of cells in the midsection along the anterior-posterior axis, as previously reported^17^, and a ventral population presenting as two bilaterally symmetric lines of cells (Fig. 3C-D and Supplementary Fig. 3E). In contrast, *H3-b* exhibited a broader expression pattern characteristic of aPSC-associated transcripts – H3-b mRNA was present in *H3-a*+ cells as well as in additional cells distributed throughout the animal in the same pattern as *piwi-1+* aPSCs (Fig. 3C-D, Supplementary Fig. 3E-F). *H3.2* and *H3.3* also exhibit similarly broad expression patterns characteristic of *piwi-1+* aPSCs (Supplementary Fig. 3G). Together, these observations indicate that *H3-a* marks spatially distinct subpopulations within the broader *H3-b*+ aPSC population.

**Figure 3.**
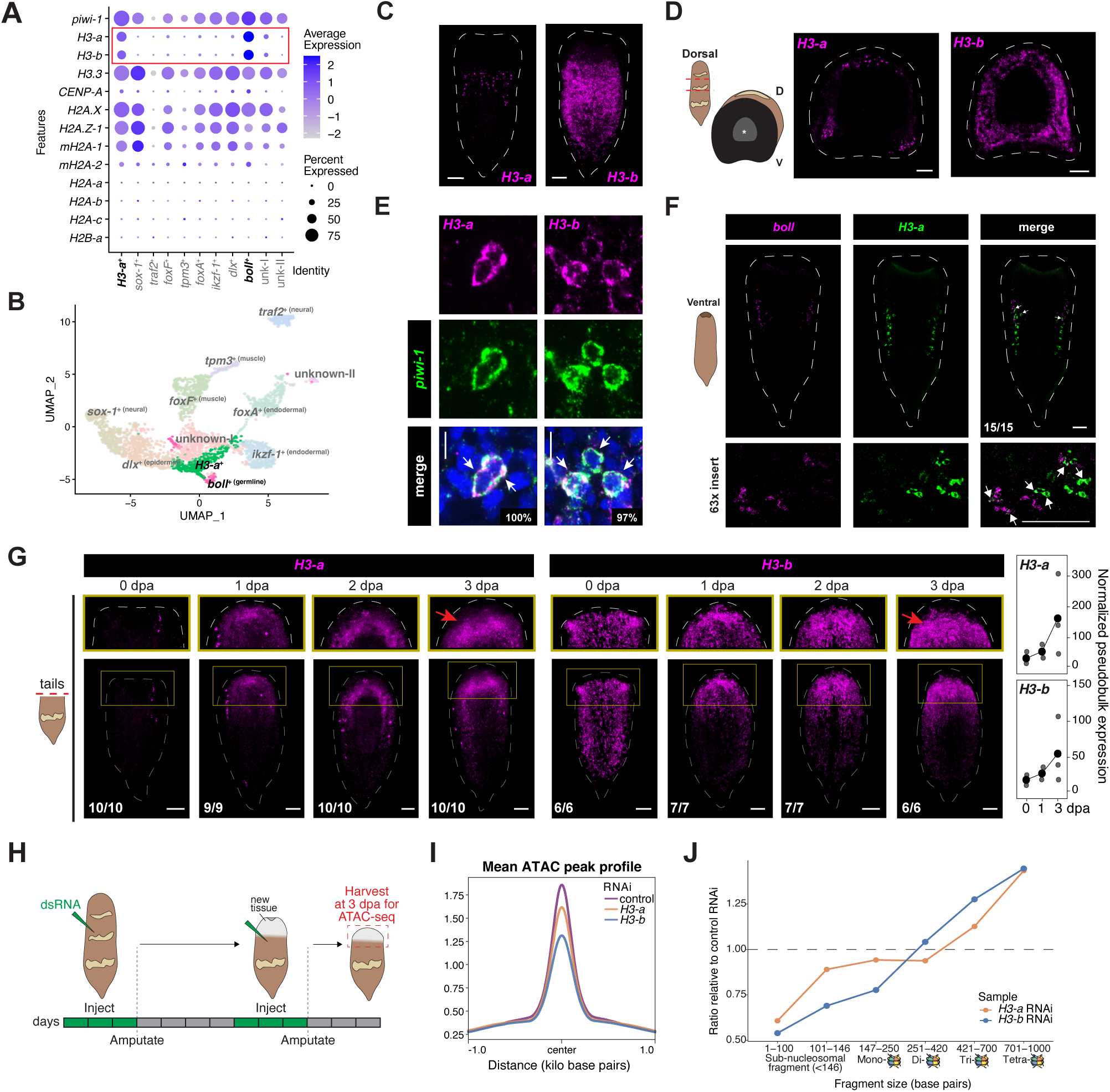
H3-a and H3-b genes are expressed in aPSCs and promote chromatin accessibility during regeneration. (A) Dot plot showing histone variant expression across *H. miamia* aPSC subpopulations defined as in Hulett et al., 2023. Dot size represents the proportion of cells expressing each histone variant, and dot color indicates average expression. *H3-a* and *H3-b* (red box) are enriched in putative unspecialized aPSCs (*H3-a*+ cluster) and *boll*+ germline-specialized aPSCs. (B) UMAP of *H. miamia* aPSC subpopulations highlighting the *H3-a*+ and *boll*+ clusters. (C) FISH for *H3-a* and *H3-b* in hatchling juveniles. Scale bars, 100 µm. (D) FISH for *H3-a* and *H3-b* in transverse views of wild-type animals following trunk amputation, highlighting dorso-ventral expression patterns. *H3-b*+ cells are broadly distributed, whereas *H3-a*+ cells occupy more spatially restricted domains. Scale bars, 100 µm; D, dorsal; V, ventral. (E) Double FISH for *H3-a* and *H3-b* with *piwi-1* in intact animals. Co-expressing cells are quantified and indicated by white arrows. Scale bars, 10 µm. (F) Ventral view of double FISH for *boll* and *H3-a*. Co-expressing cells are indicated by white arrows. Scale bars, 100 µm. (G) *Left*: Representative FISH for *H3-a* or *H3-b* in regenerating tail fragments at 0-3 days post-amputation (dpa). Histone variant expression becomes enriched near the wound by 1 dpa and becomes enriched at the base of the blastema by 3 dpa. Red arrows indicate subblastemal expression in the enlarged regions (yellow insets). Scale bars, 100 µm; numbers indicate sample size. *Right*: Normalized pseudobulk expression of *H3-a* and *H3-b* in the aPSC cluster at 0, 1, and 3 dpa, using regeneration scRNA-seq data from Hulett et al., 2023. (H) Schematic of the RNAi regime used for chromatin-accessibility profiling. Animals received dsRNA injections (green triangles) for 3 consecutive days before amputation. Tail fragments were reinjected at 5-7 dpa, amputated a second time, and collected 3 days later for ATAC-seq. Tissue harvested for ATAC-seq is indicated by the red dashed box. (I) Average ATAC-seq accessibility profile centered on called accessible regions across RNAi conditions, plotted +/− 1 kb from peak centers. (J) ATAC-seq fragment-size distributions following control, *H3-a* and *H3-b* RNAi. Ratios relative to control are plotted across fragment lengths corresponding to subnucleosomal, mono-, di-, tri-, and tetra-nucleosomal species.

Consistent with our previous observations^17^, *H3-a* and *H3-b* expression was restricted to aPSCs, with 100% of *H3-a*+ (n=444) and 97.13% of *H3-b*+ cells (n=629) co-expressing *piwi-1* (Fig. 3E). Expression of the remaining H3 family members was also enriched in aPSCs on bulk RNA-seq data (Fig. 1H). Consistently, 91.72% of *H3.2*+ (n=556) and 93.40% of *H3.3*+ (n=618) co-expressed *piwi-1* (Supplementary Fig. 3H, Supplementary File 1). In contrast, CENP-A did not exhibit significant expression within aPSC subpopulations (Fig. 3A and Supplementary Fig. 1F-G) and was therefore not examined further. As an independent assessment of aPSC-associated expression, irradiation, which eliminates dividing *piwi-1*+ cells by 7 days post irradiation (dpi)^62,63^, reduced expression of *H3-a*, *H3-b*, *H3.2 and H3.3*, indicating that all four histones are irradiation sensitive^44^ (Supplementary Fig. 3I). Together, these results confirm that *H3-a* and *H3-b*, along with the canonical *H3.2* and the variant *H3.3*, are expressed in the *H. miamia* aPSC population.

The ventral subpopulations of *H3-a*+ cells are positioned in a pattern reminiscent of the placement of female germline and developing oocytes seen in older, sexually mature *H. miamia*^17,64^. *Boll*+ cells are detected primarily among ventral *piwi-1*+ cells, with a subset co-expressing *H3-a* in hatchling and late juvenile animals (Fig. 3F and Supplementary Fig. 3J-K). Given that *boll*+ aPSCs are hypothesized to represent germline-specialized aPSCs^17^, we asked whether the ventral *H3-a*+ subpopulation also expressed markers associated with germline development in older animals. The female germ cell markers *hspb1* and *ldlr6* exhibited ventrally restricted expression in both hatchling and late juvenile animals, with a subset of these cells co-expressing *H3-a* (Supplementary Fig. 3J-K). This suggests that some ventral H3-a+ cells could be germline precursors. The male germline marker *pa1b3-2*, which expands from the ventral to the dorsal side as germ cells mature^64^, did not show co-expression with *H3-a* ventrally (Supplementary Fig. 3L). The second, dorsal *H3-a*+ subpopulation did not show expression of maturing germ cell marker genes. Thus, the ventral *H3-a*+ subpopulation co-expresses markers associated with germline precursors and female germ cells, suggesting that some, but not all, *H3-a*+ cells contribute to the germline. Notably, this association in *H. miamia* parallels the germline-restricted expression of several divergent histone variants described in other animals^3,8,11,65^. Overall, *H3-a* and *H3-b*, while co-expressed, exhibit distinct homeostatic expression patterns, with *H3-b* broadly distributed throughout the aPSC population and *H3-a* marking spatially restricted subpopulations, including a ventral population associated with germline development.

To understand the dynamics of *H3-a and H3-b* as aPSCs engage in rebuilding tissues, we followed their expression over the course of regeneration. Animals were amputated transversely to generate head and tail fragments that were collected over the course of 5 days post amputation (dpa). In tail fragments, *H3-a*+ and *H3-b*+ cells mirrored the distribution of *piwi-1*+, becoming enriched near the wound site by 1 dpa and at the base of the blastema by 3 dpa (Fig. 3G, Supplementary Fig. 3M-N). Because transverse amputations removed pre-existing *H3-a*+ cells from tail fragments, *H3-a* expression in tails corresponded to *piwi-1*+ aPSCs that upregulated *H3-a* following injury. Consistent with this observation, analysis of regeneration scRNA-seq data revealed progressively increasing *H3-a* transcript levels within aPSCs from 0 to 3 dpa (Fig. 3G, Supplementary Fig. 3N). *H3-b* transcript levels also increased progressively over the same interval, although to a lesser extent than *H3-a* (Fig. 3G, Supplementary Fig. 3N). These increases were not accompanied by upregulation of germline-associated genes, including *boll*, *hspb1*, *ldlr6* and *pa1b3-2* (Supplementary Fig. 3O), consistent with previous observations that mature germ cells decline over the same time interval during regeneration^17^. Together, these findings suggest that increased *H3-a* and *H3-b* expression in tail fragments reflects a broader regeneration-associated response within aPSCs rather than enrichment of germline-specialized aPSCs. In head fragments regenerating posterior tissues, *H3-a* and *H3-b* expression was also detected in newly formed tissue (Supplementary Fig. 3P). By 5 dpa, the expression patterns of both variants were restored to their homeostatic distributions in head and tail fragments (Supplementary Fig. 3N,P).

Together, these findings reveal distinct homeostatic expression patterns for *H3-a* and *H3-b* within aPSCs. *H3-b* is broadly expressed within aPSCs, whereas *H3-a* marks spatially restricted subpopulations, including ventral cells that may represent germline-specialized progenitors. Despite these differences, both variants are upregulated in aPSCs during regeneration, without corresponding increases in germline-associated gene expression, and become enriched at the base of the blastema, near regenerating tissues.

### H3-a and H3-b are needed for higher chromatin accessibility in regenerating *H. miamia*

Our biochemical analyses of H3-a- and H3-b-containing nucleosomes suggested that these variants could alter chromatin organization by promoting nucleosome destabilization (Fig. 2). Given that *H3-a* and *H3-b* are upregulated following amputation and enriched in aPSCs at the base of the blastema at 3 dpa in wild-type animals (Fig. 3G), we asked whether these variants contribute to global chromatin accessibility during regeneration. To capture regeneration-associated chromatin activity, we depleted either H3-a or H3-b using RNA interference (RNAi) and profiled chromatin accessibility of the anterior wound region of regenerating tail fragments at 3 dpa using ATAC-seq (Fig. 3H and Supplementary Fig. 3Q-R). Analysis of chromatin accessibility following *H3-a* and *H3-b* RNAi revealed a genome-wide shift toward reduced accessibility relative to controls (Fig. 3I, Supplementary Fig. 3S-U, Supplementary File 4).

To determine whether this reduction in accessibility was accompanied by changes in nucleosome organization, we examined ATAC-seq fragment size distributions. Both *H3-a* and *H3-b* RNAi resulted in a relative reduction of subnucleosomal fragments (DNA fragments shorter than a nucleosome) and an enrichment of longer fragments corresponding to tri- and tetra-nucleosomal fragments compared with control RNAi (Fig. 3J, Supplementary Fig. 3V, and Supplementary File 4). Together, these changes indicate that depletion of either variant results in a global reduction in chromatin accessibility in regenerative tissues. We therefore hypothesize that incorporation of H3-a and H3-b promotes a more accessible chromatin state *in vivo*.

Together with our *in vitro* assembly data (Fig. 2), these *in vivo* observations in chromatin accessibility support a model in which H3-a and H3-b generate nucleosomes that more readily permit DNA accessibility. We next sought to determine the molecular basis for these altered nucleosome properties by examining their architecture at atomic resolution.

### H3-b weakens histone-DNA contacts within the nucleosome

We used cryogenic electron microscopy (cryoEM) to investigate the structural basis for the reduced stability of nucleosomes containing H3-a or H3-b at atomic resolution. Native H3-a and H3-b nucleosomes did not remain intact under cryogenic conditions. We therefore stabilized these particles by glutaraldehyde crosslinking during sucrose gradient separation (GraFix, see Methods). Under these conditions, H3-a nucleosomes still disintegrated upon freezing on the grid, further substantiating their compromised stability observed in biochemical assays (Fig. 2G). In contrast, H3-b nucleosomes remained intact, making them amenable to single-particle cryoEM analysis to a resolution of 3.44 Å (Supplementary File 5).

The overall configuration of the histone core containing H3-b closely resembles that of the previously published structure of the human H3.3-containing nucleosome, PDB: 3AV2^66^, albeit with differences (Fig. 4A-B, Supplementary Fig. 4A). Aligning the two structures via the H3-H3’ four helix bundle reveals a high degree of conformational similarity in the histone octamer, although the H3-b-containing nucleosome is slightly more expanded along the short axis of the disc when measured between H2B ɑC helices (∼1.9 Å, Supplementary Fig. 4B). More notably, only 105 base pairs of the 601 DNA could be confidently modeled in the H3-b structure, compared to the 147 base pairs visualized in the H3.3 nucleosome crystal structure (Fig. 4B). This reduced DNA occupancy is consistent with the lower stability of H3-b nucleosomes relative to canonical nucleosomes (Fig. 2G). Finer differences at specific amino acid positions provide additional structural rationalization for this reduced stability. Two conserved arginine residues present in all H3 variants, including H3.3 (Arg64 and Arg83, in *Hofstenia miamia* H3.3 primary sequence numbering), known as sprocket arginines, insert into the DNA minor groove and stabilize histone-DNA interactions^67^. In H3-b nucleosomes, these residues are replaced by asparagine and leucine (Asn65 and Leu85, respectively), eliminating the electrostatic interactions required for stable minor groove insertion (Fig. 4C, Supplementary Fig. 4C). Similarly, a conserved lysine in H3.3 (Lys115) is replaced by Asn117 in the L2 loop of H3-b, removing two DNA contacts and reducing positive surface charge at the H3-b/H4 tetramer-dyad interface (Fig. 4D, Supplementary Fig. 4D). Together, these substitutions diminish H3-b contact points with DNA and contribute to the reduced stability of the particle.

**Figure 4.**
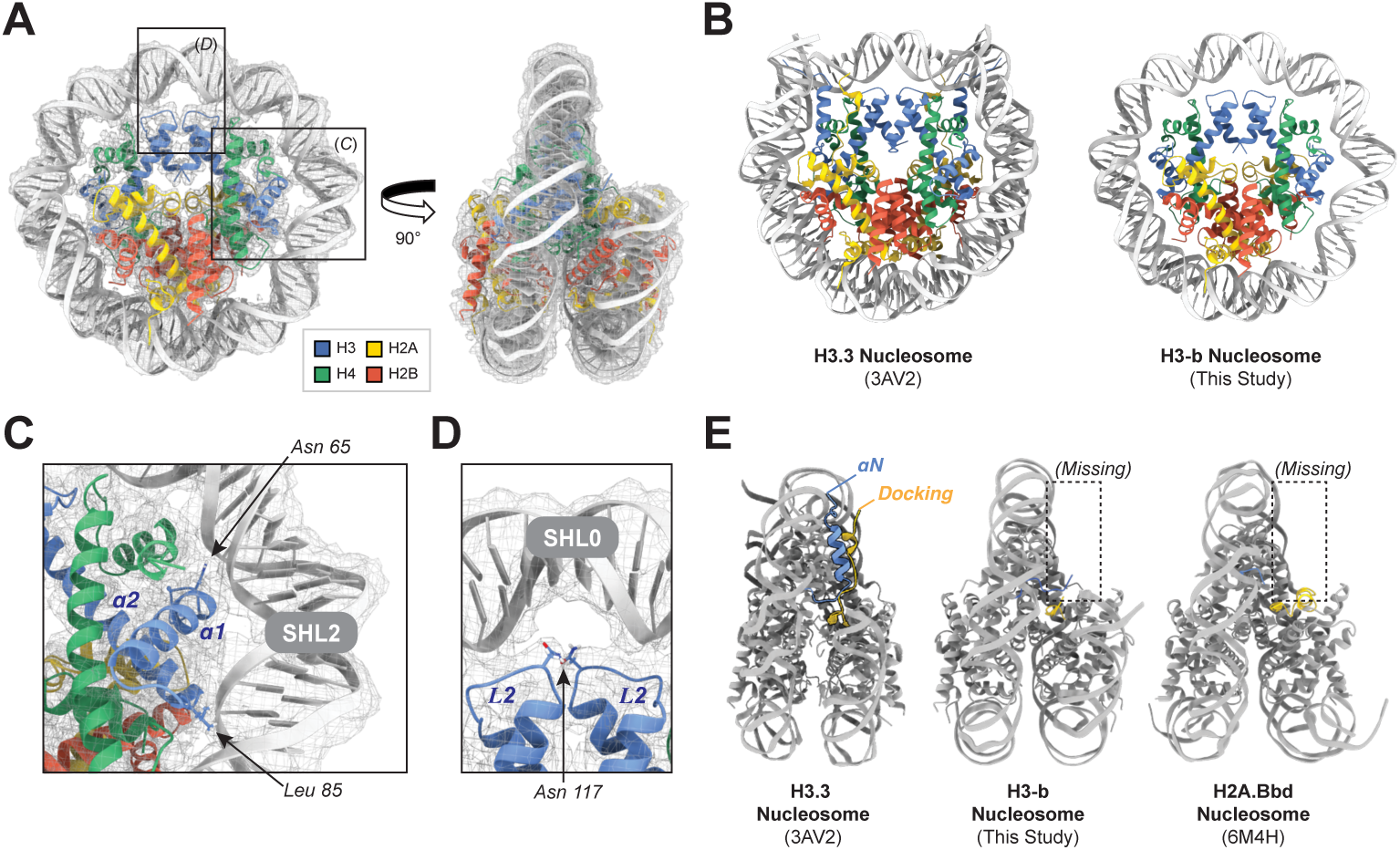
H3-b weakens histone-DNA interactions within the nucleosome. (A) Atomic model of the H3-b-containing nucleosome, assembled with human H2A, H2B, H4, and 147-bp Widom 601 DNA, superimposed with the 3.44 Å-resolution cryo-EM density map. (B) Structural comparison of the H3-b- and H3.3-containing nucleosomes (PDB: 3AV2). (C) Enlarged view of the H3-b ɑ1-loop1-ɑ2 region, highlighting the Asn65 and Leu85 near superhelical location 2 (SHL2) of nucleosomal DNA. (D) Enlarged view of the nucleosomal dyad (SHL0), showing the Asn117 sidechains from both H3-b monomers and their positions relative to DNA. (E) Side-view comparison of the H3.3-(PDB: 3AV2), H3-b- and H2A.Bbd-containing (PDB: 6M4H) nucleosomes. The H2A docking domain (yellow) and H3 ɑN helix (blue), or their corresponding disordered/absent regions, are highlighted by the dotted box. Histone proteins are color-coded: H3, blue; H4, green; H2A, yellow; H2B, red.

In addition to these localized changes, the first 57 residues of H3-b (spanning the N-terminal tail through the end of the ɑN helix) could not be modeled into the density map, suggesting pronounced conformational flexibility in this region (Fig. 4E, Supplementary Fig. 4E). Pairwise alignment of the N-terminal sequences of H3-b and human H3.3 demonstrates extensive divergence in a region of the NTT normally sandwiched between the exit and entry DNA gyres. In particular, two arginine residues that contact the DNA minor groove in H3.3 (Arg40 and Arg42) are replaced by histidine and glutamine in H3-b (His41 and Gln43), respectively (Supplementary Fig. 4E). These substitutions may weaken conserved histone-DNA interactions and contribute to the lack of ordered density observed in the adjacent ɑN region of H3-b. Interestingly, H3-a also contains His41 but has a positively charged lysine at position 43 (Supplementary Fig. 2M), suggesting that both divergent variants may alter histone-DNA interactions within this region through distinct electrostatic properties.

We also observed a loss of ordered density in the final 20 residues of H2A, comprising the C-terminal ‘docking domain’. Perturbations of the H2A docking domain can compromise nucleosome stability, as exemplified by the human H2A.Bbd variant, which lacks the terminal ∼20 residues of the docking domain and C-terminal tail^68,69^. Comparing our structure with the published model of the H2A.Bbd nucleosome (PDB: 6M4H)^70^, we find a similar disruption of the H2A-H3 interface, despite the fact that both structures contain entirely distinct histone variants (Fig. 4E). Because our H3-b-containing nucleosomes contain canonical H2A with an intact docking domain, the loss of ordered density in this region suggests that sequence divergence within H3-b may disrupt interactions that normally stabilize the H2A docking domain. Perturbation of this interface may further contribute to the reduced stability and DNA wrapping observed in the H3-b-containing nucleosome^68–72^.

Overall, our structural analyses demonstrate that H3-b-containing nucleosomes preserve the general architecture of the nucleosome while exhibiting weakened histone-DNA interactions and reduced DNA wrapping. The loss of conserved DNA-contacting residues, together with increased structural disorder near the DNA entry and exit regions and the H2A docking domain, provides a mechanistic explanation for the reduced stability of H3-b-containing nucleosomes and suggests a potential basis for the contribution to chromatin accessibility *in vivo*.

### H3-a and H3-b represent an understudied landscape of divergent animal H3 proteins

The structural similarity between nucleosomes containing the divergent H3-b or H2A.Bbd proteins suggests convergent evolution toward similar structural and functional outcomes. This raised the broader question of whether the divergent H3 proteins identified in *H. miamia* are outliers within the evolutionary landscape of animal H3 proteins. To investigate this, we surveyed previously deposited histone H3 sequences across animal phyla to characterize the extent of sequence divergence and whether it preferentially affects the regulatory NTT or the structural HFD. Using the human canonical H3.1 (NP_003520.1) as a query, we performed NCBI BLAST searches to retrieve putative H3 homologs across metazoans (see Methods). The recovered sequences clustered together with known H3 sequences in t-SNE space, separately from clusters containing H2A, H2B, and H4 sequences, supporting their classification as H3-like proteins (Supplementary Fig. 5A).

The NTT and HFD boundaries were defined for each sequence using homology-based domain annotation with HMMER and InterProScan (see Methods). A paired comparison of NTT and HFD percent identity, using the 10^th^ percentile of each distribution as a threshold, partitioned H3 sequences into four distinct classes (Fig. 5A, Supplementary Fig. 5B, Supplementary File 6). The vast majority of H3 sequences fell into the canonical-like class (upper-right, Fig. 5A), with high identity in both domains (1235/1430, 86.4%). Previous studies have characterized histone H3 variants that, despite harboring amino acid substitutions, remain closely related to established variants such as H3.3 and retain canonical-like nucleosome properties^3–6,10^. The remaining three classes instead captured different modes of H3 divergence: conserved HFD with divergent NTT (upper-left), conserved NTT with divergent HFD (lower-right), or divergence in both domains (lower-left). Notably, *H. miamia* H3-a and H3-b occupy this latter class where both HFD and NTT domains are divergent (Fig. 5B).

**Figure 5.**
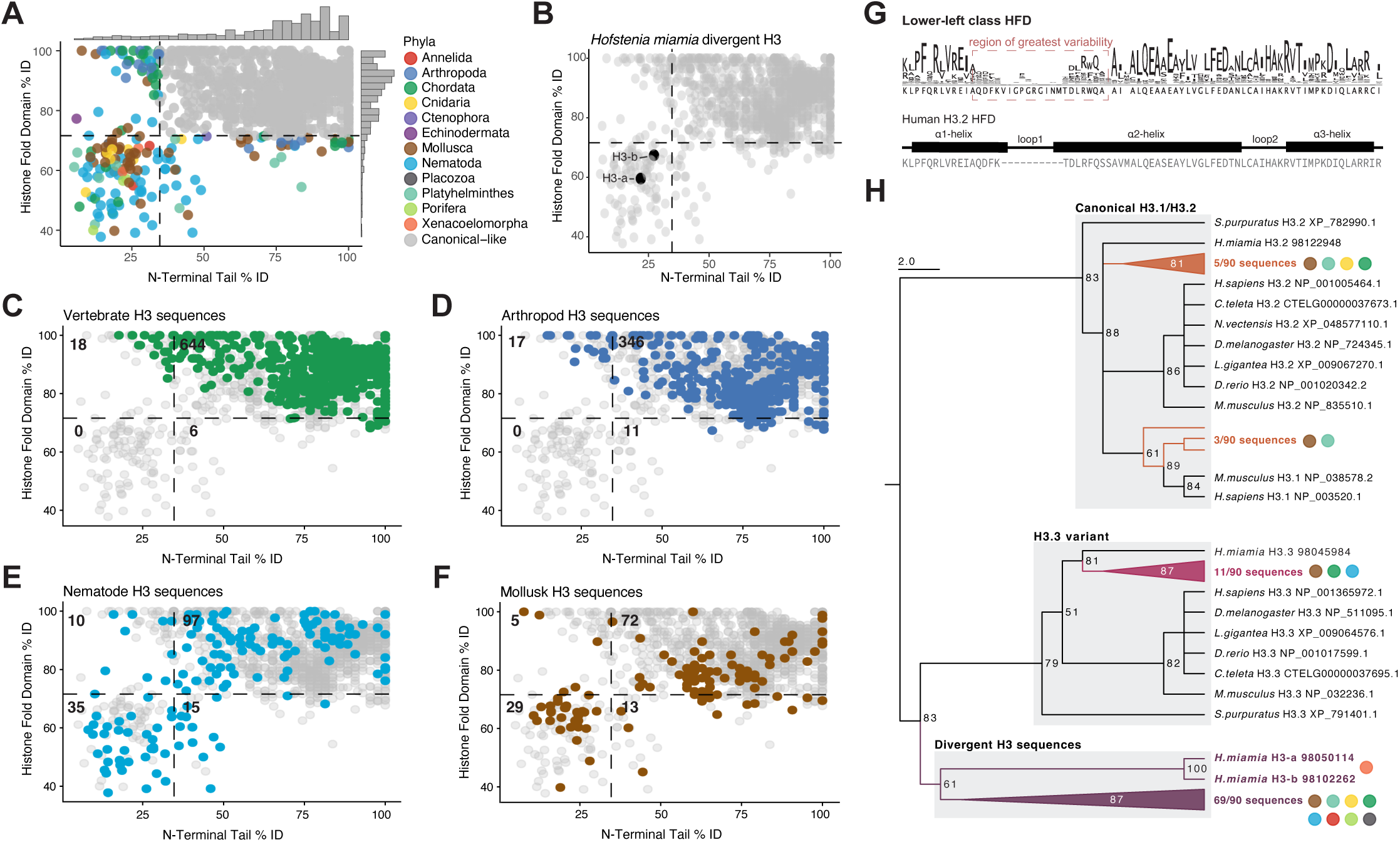
Divergent histone H3 proteins occur across animal phyla and reflect multiple evolutionary origins. (A) Paired comparison of N-terminal tail (NTT) and histone fold domain (HFD) percent identity among recovered animal H3 sequences. Each point represents one H3 sequence and is colored by phylum. Dashed lines indicate the 10th percentile of the NTT and HFD identity distributions, partitioning sequences into four divergence classes. Marginal histograms show the distribution of sequence identity along each axis. (B) Position of *H. miamia* H3-a and H3-b within the lower-left class, characterized by divergence in both the NTT and HFD. (C-F) Distribution of vertebrate (C), arthropod (D), nematode (E), and molluscan (F) H3 sequences across divergence classes. Sequences from the indicated groups are colored, with all other H3 sequences shown in gray. Numbers indicate sequence counts within each quadrant. (G) HFD sequence conservation among lower-left class H3 proteins compared with human H3.2. The sequence logo shows residue frequency at each aligned position, with letter height proportional to residue frequency. The region of greatest sequence variability is indicated above the logo. The structure of the human H3.2 HFD is shown below for reference. (H) Maximum-likelihood phylogenetic tree of lower-left class H3 sequences together with representative canonical H3.1/H3.2 and H3.3 proteins. Lower-left sequences are shown as collapsed clades: 8 of 90 associate with H3.1/H3.2 lineage, 11 of 90 with the H3.3 lineage, and the remaining 71 form a divergent lineage outside these canonical clades that includes *H. miamia* H3-a and H3-b. Colored circles indicate the animal phyla represented within each collapsed clade; node values indicate bootstrap support.

Having categorized the recovered H3 sequences, we next examined how the divergent classes were distributed across animal phyla. The most abundantly sampled phyla were chordates and arthropods. Among chordate sequences, 97.4% (n=668) were derived from vertebrates and 2.6% (n=18) from invertebrate chordates (Fig. 5C, Supplementary Fig. 5B). Vertebrate H3 sequences were overwhelmingly canonical-like, with 644 sequences (96.4%) in the upper-right class. Among the divergent vertebrate H3s, sequences with a divergent NTT but conserved HFD (upper-left: n=18) were more common than sequences with a divergent HFD but conserved NTT (lower-right: n=6). This pattern is consistent with previously characterized chordate H3 variants that exhibit substantial NTT divergence while retaining a conserved, nucleosome-competent HFD^65,73–75^. Notably, no divergent vertebrate H3 occupied the lower-left class; instead, all six chordate sequences in this class were derived from invertebrate chordates (Supplementary Fig. 5B). Similarly, arthropod sequences were entirely absent from the lower-left (Fig. 5D), suggesting that in our dataset, arthropod H3 sequences diverge in either the NTT or the HFD, but not both.

In contrast, HFD divergence was substantial among nematode and molluscan H3 sequences (Fig. 5E-F). Together, these two phyla accounted for 64 of the 90 sequences in the lower-left class and 28 of the 52 sequences in the lower-right class. Relative to their representation in the overall dataset, nematodes were enriched 3.5-fold and 2.6-fold in the lower-left and lower-right classes, respectively, while mollusks were enriched 3.9-fold and 3.0-fold (Supplementary File 6). Divergence class was strongly associated with phylum overall (Chi-squared = 375.9, p < 10^−4^), driven primarily by an excess of chordate sequences in the canonical-like upper-right class and of nematode and molluscan sequences in the HFD-divergent lower classes (Supplementary File 6). Although nematodes and mollusks accounted for most sequences in the lower-left class, this pattern of extensive H3 divergence was also observed in additional animal phyla (Supplementary Fig. 5B). Further, no H3 sequences from ctenophores or placozoans in this dataset were assigned to the canonical-like class (Supplementary Fig. 5B).

Given the substantial divergence of sequences in the lower-left class, we next asked whether these sequences retain recognizable H3 structural features and how this divergent state evolved across animal lineages. Despite their overall divergence, lower-left H3 sequences retained recognizable H3 sequence organization across much of the HFD, with the greatest sequence variability concentrated within the ɑ1-loop1-ɑ2 region (Fig. 5G). AlphaFold2 predictions further indicated that these divergent HFDs retain the canonical histone-fold conformation (Supplementary Fig. 5C). Phylogenetic analysis placed subsets of lower-left sequences within both the canonical H3.2 and H3.3 lineages, whereas a larger group, which includes *H. miamia* H3-a and H3-b, formed a distinct lineage outside these clades (Fig. 5H, Supplementary Fig. 5D). Thus, the lower-left class consists of highly divergent H3 sequences that appear to reflect at least three independent evolutionary origins: one within the H3.2 lineage, one within the H3.3 lineage, and a third outside these canonical clades that includes H3-a and H3-b.

Together, these observations reveal a broader evolutionary landscape of animal H3 proteins, in which sequence divergence encompasses both the regulatory NTT and structural HFD across multiple animal lineages.

## DISCUSSION

Although histones are among the most conserved eukaryotic proteins, their conservation coexists with considerable sequence diversity among histone variants. While the functions of many conserved variants are well-established, the molecular properties of more extensively divergent metazoan histones remain comparatively poorly understood. Here, we characterized two *H. miamia*-specific H3 paralogs, H3-a and H3-b, that assemble into nucleosomes with reduced stability and incomplete DNA wrapping, and contribute to chromatin accessibility during whole-body regeneration. Their unusual molecular properties, together with their enrichment in aPSCs, raise questions about how histone sequence diversification influences chromatin regulation and cellular identity.

Our cross-phyla survey placed H3-a and H3-b within a broader repertoire of highly divergent H3 proteins represented across multiple animal lineages. Phylogenetic analysis suggests that this extensive divergence arose through multiple evolutionary trajectories, including within the established H3.2 and H3.3 lineages and in a separate group containing H3-a and H3-b (Fig. 5). Within our dataset, chordates and arthropods, which include well-studied model organisms, showed the lowest representation of divergent H3 classes, whereas several less-studied phyla contributed the majority of divergent sequences. The enrichment observed in nematodes is consistent with previous studies documenting divergent H3.3 paralogs^3,10^, as well as the elevated rates of sequence evolution characteristic of this phylum^76^. However, the comparable enrichment in mollusks, whose genomes evolve relatively slowly^77^, suggests that extensive histone H3 diversification may not be explained solely by higher rates of sequence evolution. Our characterization of H3-a and H3-b provides a starting point for investigating the molecular properties of highly divergent H3 proteins across animal lineages. Beyond H3-a and H3-b, our annotation of the *H. miamia* histone repertoire also identified divergent H2A and H2B proteins that fall outside conventional variant classifications (Supplementary Fig. 1A). Although not characterized further here, these variants provide additional opportunities to investigate how histone sequence diversification influences nucleosome properties and chromatin regulation.

How divergent histones alter nucleosome structure and stability is central to understanding their function. Despite their extensive sequence divergence, both H3-a and H3-b were capable of assembling nucleosomes *in vitro*, but these particles exhibit reduced stability and incomplete DNA wrapping. The H3-b cryoEM structure provides a mechanistic explanation for these properties: while the overall histone core is preserved, specific substitutions weaken interactions that normally stabilize nucleosomal DNA (Fig. 4), providing a route by which sequence divergence can increase nucleosome dynamics^54^. Disorder in the H2A docking domain, leading to weak H3-H2A and histone-DNA interactions, is a hallmark of nucleosomes containing the divergent human H2A variant H2A.Bbd^68–72^. In our case, however, a comparable outcome is achieved by divergent sequence characteristics in the H3-b ɑN region that abrogate its interaction with and stabilization of the H2A docking domain (Fig. 4E). Together, these observations suggest that nucleosome destabilization and enhanced DNA accessibility can arise convergently through distinct structural perturbations of the histone core. Given that many of the highly divergent H3 proteins recovered in our survey retain predicted histone-fold architecture (Fig. 5G), future biochemical and structural characterization will be essential to determine whether they exhibit similar nucleosome properties or have evolved distinct mechanisms of chromatin regulation.

Our data showed that global chromatin accessibility is reduced following *H3-a* or *H3-b*depletion, suggesting that these variants contribute to maintaining a permissive chromatin environment during regeneration. Work in planarians has identified a permissive chromatin environment associated with stemness genes in aPSCs, supported by intrinsic DNA sequence properties with low nucleosome affinity^78^. Although this work suggests a different underlying cause for chromatin accessibility than our study, together these findings raise the possibility that broad chromatin accessibility may be important for aPSC function in two evolutionarily distinct regenerative systems. The enrichment of *H3-a* and *H3-b* in aPSCs raises the further interesting question of whether their influence on chromatin accessibility contributes to stem cell potency. In vertebrate embryonic stem cells, conserved histone variants H3.3 and H2A.Z contribute to chromatin organization at regulatory elements associated with pluripotency and lineage commitment^21,26,79^. Whether the chromatin changes observed following H3-a and H3-b depletion directly influence aPSC potency remains to be resolved, but their regeneration-responsive upregulation and role in chromatin organization *in vivo* (Fig. 3) are nevertheless consistent with a model in which these variants help maintain a pluripotent state in aPSCs. More functional work in *H. miamia* and other species with aPSC-like populations, such as planarians and cnidarians, is needed to fully elucidate this relationship and assess whether histone variants in other species contribute to similar regulatory processes. Given the similarities between aPSCs across species^15^, future studies could investigate the extent to which changes in the histone repertoire enable aPSC function across animals.

In addition to their roles in aPSCs, *H. miamia* divergent histones also likely impact germline biology. Our observations support a germline-associated role for the ventral *H3-a*+ subpopulation, whereas the dorsal *H3-a*+ subpopulation likely corresponds to an unspecialized or transient aPSC subpopulation^17^. The association of a highly divergent histone with the germline is particularly intriguing given that several previously characterized histone variants with specialized functions also exhibit germline-enriched or testis-specific expression^3,8,11,65^. This recurring germline association raises the possibility that H3-a and H3-b originally evolved for germline-specific functions, as have variants in other species, and then later evolved functions in stem cell-mediated regeneration. The close relationship between aPSCs and the germline has been recognized for decades, as many components of the germline multipotency module are expressed in aPSCs across species^14^.

Together, our findings establish H3-a and H3-b as *H. miamia*-specific histone H3 variants that escape canonical chaperone recognition, destabilize nucleosomes, and promote chromatin accessibility in the context of whole-body regeneration. More broadly, the characterization of H3-a and H3-b provides molecular insight into highly divergent histone proteins represented across animal lineages, while positioning *H. miamia* as a tractable system for investigating how histone sequence diversification may contribute to chromatin regulation and stem cell biology.

## MATERIALS AND METHODS

### Identification of histone and chaperone genes

Curated protein sequences corresponding to nucleosomal histone families (H2A, H2B, H3 and H4) and their respective variants were obtained from the NCBI Histone Database 2.0^30^. These sequences were used as queries to identify homologous genes in *Hofstenia miamia* through BLAST searches against the genome assembly^31^ and transcriptome^80^. Genomic loci corresponding to identified histone genes were annotated based on sequence similarity and visualized with TBtools^81^. Three-dimensional protein structures were predicted using AlphaFold2^82^. Putative 3’ untranslated region (UTR) stem-loop structures were predicted using R-scape^38^ and Rfam^37^. Putative histone chaperone homologs were identified using a similar approach, employing curated protein sequences of known chaperones as queries.

### Phylogenetic analysis

Protein sequences were aligned using MAFFT v.7.490^83^ with the BLOSUM62 scoring matrix and default gap opening penalties. Phylogenetic trees were inferred using IQ-TREE v2.3.5^84^. The best-fitting amino acid substitution model was selected using ModelFinder (-m MFP), allowing estimation of empirical amino acid frequencies, invariant sites, and rate heterogeneity where appropriate. Branch support was assessed using 1,000 ultrafast bootstrap replicates^85^. Trees were visualized in FigTree v.1.4.4.

### Single-cell RNA-seq analysis

Single-cell RNA sequencing data from *H. miamia* hatchling juvenile and the merged regeneration comprising post-amputation timepoints were reanalyzed using the same clusters validated by^17^. UMAP embeddings and DotPlots of genes of interest were generated using DimPlot and DotPlot functions from Seurat v5.0.0^86^. The merged regeneration dataset was subset to the 0, 24, and 72 hpa timepoints. Each timepoint replicate was aggregated into pseudobulk profiles using AggregateExpression. The resulting count matrix was analyzed using DESeq2 v.1.5^87^, and pairwise contrasts were performed between 0 vs 24, and 0 vs 72 hpa. P-values were adjusted for multiple testing using the Benjamini-Hochberg procedure using default parameters. Claude Sonnet 5 was used to streamline data processing and visualization.

### Bulk RNA-seq analysis

Bulk RNA sequencing data of Hoechst^high^ and Hoechst^low^, as well as irradiated and unirradiated *H. miamia* samples were reanalyzed using the same parameters published in^44^. Fold changes in expression for genes of interest were calculated as log_2_ of the ratio of the mean transcript per million reads (TPM) values across replicates between conditions. Volcano plots were generated using Sleuth^88^ and expression heatmaps were generated using pheatmap. Claude Sonnet 5 was used to streamline data processing and visualization.

### Animal maintenance

Juvenile and adult *Hofstenia miamia* worms were maintained in artificial seawater made and filtered with Instant Ocean Sea Salt (37 ppt, pH = 7.8-8.0) at 21 °C. Juvenile animals were grown in zebra fish tanks, while adult animals were maintained in plastic boxes. Seawater was replaced twice a week, followed by feeding rotifers *Brachionus plicatilis* to juvenile worms, while feeding freshly hatched brine shrimp *Artemia* sp to adult worms. Embryos were collected using a glass pipette twice a week prior to seawater replacement.

### Histone protein extraction

Animals were dissociated as previously described^17^, yielding cell suspensions in calcium- and magnesium-free artificial sea water (CMF-ASW). Cells were pelleted and resuspended in a hypotonic lysis buffer (10 mM Tris-HCl at pH 8.0, 1 mM KCl, 1.5 mM MgCl2) and incubated for 30 minutes at 4 °C by rocking in a nutator to promote cell swelling and membrane rupture, enabling nuclear isolation^89^. Isolated nuclei were resuspended in 0.2 M sulfuric acid and incubated overnight at 4 °C to extract histone proteins^90^. Samples were centrifuged for 20 minutes at 12,000 rcf at 4 °C to remove insoluble material, and histone proteins were precipitated by addition of trichloroacetic acid at a final concentration of 12% m/v and incubated over ice for 2 hours^91^. Samples were centrifuged and pellets were washed with acetone and resuspended in molecular-grade water. Protein concentrations were determined using the Bradford assay (Bio-Rad), with absorbance measured at 595 nm using a NanoDrop spectrophotometer. Standard curves were generated using bovine serum albumin (Thermo Fisher, #23209).

### Mass spectrometry sample preparation and analysis

Histone samples were prepared in an SDS-containing buffer with DTT at a final concentration of 1 mM and separated by SDS-PAGE using 15% separation gels. Gel regions corresponding to 10-15 kDa were excised using a sterile razor blade and submitted to The Taplin Biological Mass Spectrometry Facility (Harvard Medical School) for LC-MS/MS analysis. Excised bands were cut into approximately 1 mm^3^ pieces and subjected to a modified in-gel trypsin digestion procedure^92^. Gel pieces were washed, dehydrated with acetonitrile, dried in a speed-vac, and rehydrated at 4 °C with 50 mM ammonium bicarbonate containing 12.5 ng/µL modified sequencing-grade trypsin (Promega, Madison, WI). After 45 minutes, excess trypsin solution was removed and replaced with 50 mM ammonium bicarbonate sufficient to cover the gel pieces, and samples were incubated overnight at 37 °C. Peptides were extracted with a solution containing 50% v/v acetonitrile and 1% v/v formic acid, dried in speed-vac for 1 hour, and stored at 4 °C until analysis. On the day of the analysis, samples were reconstituted in 5-10 µL of HPLC solvent A (2.5% v/v acetonitrile, 0.1% v/v formic acid). A nano-scale reverse-phase HPLC capillary column was created by packing 2.6 µm C18 spherical silica beads into a fused silica capillary (100 µm inner diameter x 30 cm length) with a flame-drawn tip^93^. After equilibrating the column, each sample was loaded via a Thermo EASY-LC (Thermo Fisher Scientific, Waltham, MA). Peptides were eluted with an increasing gradient of HPLC solvent B (90% v/v acetonitrile, 0.1% v/v formic acid), ionized by electrospray, and analyzed on an Orbitrap Exploris 480 mass spectrometer (Thermo Fisher Scientific, Waltham, MA). Fragmentation spectra were searched using Sequest^94^ (Thermo Fisher Scientific) against a custom database of *H. miamia* histone proteins. To assess specificity, non-histone nuclear proteins within a similar molecular weight range (98009552-LSM2 and 98057746-LSM3) were included in the database as negative controls. All databases included a reversed version of all sequences, and data were filtered to a peptide false discovery rate of 1-2%. The percentage of the total detected peptides was calculated on a square root scale and plotted to visualize abundances.

### Fixation and *in situ* hybridization

Animals were fixed in 4% v/v formaldehyde in 1x PBS containing 0.1% Triton X-100 at room temperature for 1 hour by rocking on a nutator. Samples were dehydrated in 100% methanol and stored at −20 °C until use. Riboprobes labeled with digoxigenin or fluorescein were synthesized as previously described^62^. Fluorescent *in situ* hybridization (FISH) was performed following established protocols^63^ with the following modification: Proteinase K digestion was extended for 20 minutes on larger animals, and development solutions were prepared using the respective riboprobe at a 1:1000 dilution.

### Irradiation experiments

Animals were exposed to a cesium irradiation source at a dose of 10,000 rads. Following irradiation, animals were maintained for 7 days to allow depletion of proliferative cells^62^. Animals were either fixed at 7 days post irradiation or amputated and allowed to regenerate for subsequent time points prior to fixation.

### Amputation and regeneration assays

Animals were amputated transversely along the middle stripe using a razor blade to generate head and tail fragments. Fragments were allowed to regenerate for the determined amount of days prior to fixation or downstream processing. Regenerative outcomes were assessed based on blastema formation and/or the expression of known tissue markers in the new tissue by *in situ* hybridization.

### Amplicon-sequencing and analysis

Primers were designed to amplify regions shared between the H3-a and H3-b genes (forward: 5’-CCCCTCGAAAAAGTATGCAACC-3’; reverse: 5’-TTAAATGGCTTGACCACCGC-3’). PCR amplification was performed using plasmid templates containing the H3-a or H3-b target sequences. Reactions were carried out using 2x ES Taq Master (CWBio, #CW0690H) mix under the following cycling conditions: initial denaturation at 94 °C for 2 minutes; 40 cycles of denaturation at 94 °C for 25 seconds, annealing at 59.5 °C for 40 seconds, and extension at 72°C for 1 minute; followed by a final extension at 72 °C for 2 minutes. PCR products were purified using the QIAquick PCR and Gel Cleanup kit (QIAGEN, #28104) according to the manufacturer’s instructions and quantified using a Qubit fluorometer. Purified amplicons were submitted to Genewiz for Amplicon-EZ sequencing at a concentration of 20 ng/µL. Sequencing reads were mapped to reference sequences using Bowtie 2^95^, and read counts were used to quantify the relative representation of each target gene.

### Lentiviral expression and imaging of histone-GFP fusion proteins

*Hofstenia miamia* histone proteins were synthesized as GeneBlocks through Integrated DNA Technologies and cloned into a lentiviral backbone under the control of a tet-ON promoter and fused to a C-terminal GFP^54^. The lentiviral backbone used additionally contained a G418 resistance cassette. Lentivirus was prepped by transfecting HEK293T LentiX cells, and filtered media was added to NIH-3T3 cell media (DMEM containing 10% v/v FBS and 1% m/v Penicillin/Streptomycin). After 48 hours, NIH-3T3 cell media was replaced with fresh media containing 500 µg/mL G418, and cells were allowed to undergo selection for 5 days. For imaging experiments, cells were treated with doxycycline (DOX) for 24 hours, and then with either DMSO (control) or colcemid overnight (∼16 hours) in DOX-containing media. Cells were treated with CellMask near-infrared for 10 minutes followed by fixation as per manufacturer’s instructions, and stained with Hoechst in PBS for 10 minutes. Following 3x PBS washes, cells were imaged on a Nikon Ti2 Eclipse microscope equipped with the ANDOR Dragonfly confocal system. Segmentation and quantification of images were performed using Fiji^96^ and results were visualized with ggplot2 v.3.

### Expression and purification of recombinant histone proteins

Histone gene constructs were synthesized using the Flash Gene synthesis service and subsequently cloned into expression vectors by GenScript. Recombinant histones were expressed and purified following established protocols developed by Pamela N. Dyer and colleagues with minor modifications^56^. Individual histone genes were transformed into *E. coli* CodonPlus RIPL cells (Agilent) and grown in LB medium supplemented with appropriate antibiotics. Cultures were grown at 37 °C to an OD₆₀₀ of ∼0.6 and induced with IPTG for 3 h. Cells were harvested by centrifugation and either stored at −80 °C or processed immediately. Cell pellets were resuspended in a wash buffer (50 mM Tris-HCl pH 7.5, 100 mM NaCl) and lysed by sonication. Histones were recovered from inclusion bodies, which were isolated by centrifugation. The insoluble fraction was subjected to multiple washing steps using the wash buffer supplemented with 1% (v/v) Triton X-100 to remove contaminating proteins, lipids, and nucleic acids. Purified inclusion bodies were solubilized in 25 mL of unfolding buffer (50 mM Tris-HCl pH 7.5, 7 M guanidine hydrochloride, 5 mM β-mercaptoethanol). The solubilized material was clarified by centrifugation and loaded onto a Sepharose 200 size-exclusion column (column volume, 1750 mL) equilibrated in denaturing buffer (25 mM Tris-HCl pH 7.5, 7 M urea, 200 mM NaCl, 5 mM β-mercaptoethanol). Fractions containing histones were pooled and further purified by cation-exchange chromatography (SP Sepharose, column volume, 30 mL) under denaturing conditions using the same buffer system, with histones eluted over a linear gradient to 1 M NaCl. Fractions containing the target histone were identified by SDS-PAGE, pooled, and dialyzed extensively against water supplemented with a reducing agent (β-mercaptoethanol or DTT) to remove denaturant and salt. The dialyzed histones were subsequently lyophilized and stored at −20 °C until further use. The molecular weight of the purified proteins was confirmed by MALDI–TOF/TOF mass spectrometry using a Bruker UltrafleXtreme instrument at the Analytical Resources Core at Colorado State University.

### Histone Purification and Dimer/Tetramer Assembly

Plasmids containing the open reading frames for *H. miamia H3-a* and *H3-b* were submitted to the Histone Source at Colorado State University for expression and purification. Lyophilized aliquots of purified H3-a and H3-b were dissolved in the unfolding (UF) buffer (20 mM Tris, 6 M guanidinium chloride, pH 7.5) to final concentrations of 1 mM. To assemble (H3-H4)_2_ tetramers, each H3 variant (including the canonical H3.2 from *H. sapiens*, also purified by the Histone Source) was mixed 1:1 with *H. sapiens* H4 (The Histone Source) in UF buffer to final concentrations of 70 µM in a 1 mL total volume. These mixtures were transferred to D-Tube Maxi dialysis cassettes (Millipore Sigma) outfitted with dialysis membranes (6-8 kDa molecular weight cutoff (MWCO)) and suspended in 400mL pre-chilled refolding high salt buffer (‘RB High’: 10 mM Tris, 1.75 M NaCl, 0.25 M L-arginine, 5 mM TCEP, 0.25 mM EDTA, pH 7.5). Dialysis was allowed to proceed overnight with gentle stirring at 4°C. Dialysate was removed at a constant flow rate of 1.5 mL min^−1^ while being replenished with fresh RB High at the same rate. Reconstituted histone complexes were resolved on a Superdex 200 10/300 GL column (Cytiva Life Sciences) pre-equilibrated in RB High. Elution was performed at a constant flow rate of 0.3 mL min^−1^ at 4 °C on an AKTA Purifier system (Cytiva). Collected fractions were analyzed by SDS-PAGE, pooled, and concentrated in Amicon Ultra-15 centrifugal concentration units (10 kDa MWCO membrane; EMD Millipore). (H3-H4)_2_ tetramers were stored at 4°C in RB High at concentrations ranging from 30-50 µM. The H2A-H2B dimer was prepared and handled in the same manner, except with *H. sapiens* H2A and H2B (The Histone Source) provided as inputs. Dimer stocks were concentrated and stored similarly in RB High at concentrations of 50-80 µM.

### Nucleosome Reconstitution

A 147 base pair double stranded Widom 601 DNA sequence (‘147DNA’) was used in all nucleosome reconstitution reactions: ATCTGAGAATCCGGTGCCGAGGCCGCTCAATTGGTCGTAGACAGCTCTAGCACCGCTTAA ACGCACGTACGCGCTGTCCCCCGCGTTTTAACCGCCAAGGGGATTACTCCCTAGTCTCCA GGCACGTGTCAGATATATACATCCGAT. All components (DNA, (H3(.2 or -a or -b)-H4)_2_ tetramer, H2A-H2B dimer) were exchanged into RB High (10 mM Tris, 1.75 M NaCl, 0.25 M L-arginine, 5 mM TCEP, 0.25 mM EDTA, pH 7.5) if not already present in that buffer. 45 µL reconstitution reactions were prepared on ice, containing 3 µM 147DNA and empirically-determined optimal concentrations of histone tetramer and dimer. For H3-a containing samples, a 1:1.3:2.86 DNA:tetramer:dimer ratio was used; for H3.2 and H3-b containing samples, a 1:1.4:3.08 DNA:tetramer:dimer was used. In samples lacking H2A-H2B, the corresponding volume of dimer was replaced with RB High. Reconstitution mixtures were transferred to 50 µL dialysis cassettes that were affixed with small squares of pre-rinsed 6-8 kDa MWCO regenerated cellulose dialysis membrane (Spectrapor). Cassettes were then suspended in 400 mL pre-chilled RB High and allowed to dialyze overnight with rapid stirring at 4 °C. During dialysis, RB High from the dialysate was removed using a peristaltic pump at a constant rate of 1.5 mL min^−1^, while simultaneously being replenished with refolding buffer 150, ‘RB150’ (10 mM Tris, 150 mM NaCl, 5 mM TCEP, 0.25 mM EDTA, pH 7.5), at the same flow rate. The following day, the cassettes were transferred to another beaker containing 500 mL refolding buffer 2.5, ‘RB2.5’ (10 mM Tris, 2.5 mM NaCl, 5 mM TCEP, 0.25 mM EDTA, pH 7.5), and allowed to continue dialyzing for another 16-18 hours at 4°C. Following this dialysis, reconstitution mixtures were removed from the dialysis cassettes and transferred to 1.5 mL Eppendorf tubes, sedimented at maximum speed for 5 minutes at 4°C in a microcentrifuge, and analyzed by native gel electrophoresis. All nucleosome samples were diluted in 12% w/v Ficoll 400 and loaded into the wells of a 5% polyacrylamide tris-borate-EDTA (TBE) gel. Electrophoresis was performed at a constant voltage (90 V) for 60 minutes with 0.2x TBE (17.8 mM Tris, 17.8 mM sodium borate, 0.4 mM EDTA, pH 8.0) used for the running buffer. Gels were imaged via ethidium bromide staining on an Azure 600 imaging system (Azure Biosystems). Nucleosome reconstitutions for biophysical and TEM experiments were performed at a 5x scale, with all volumes scaled accordingly. Concentrations post-reconstitution were measured via 254 nm absorbance on a Nanodrop One^C^ spectrophotometer (Thermo Scientific), using the characteristic absorbance ratio of 0.531 mM AU^−1^ calculated for 147DNA (assuming one double stranded 147DNA duplex per nucleosome particle). Nucleosome samples were stored in RB2.5 at 4 °C until further use.

### Mass Photometry

All mass photometry data were collected on a Two^MP^ Mass Photometer (Refeyn Ltd.) outfitted with a 488 nm laser. Prior to data acquisition, High Precision glass coverslips (thickness no. 1.5H, Marienfeld-Superior) were repeatedly rinsed with water and isopropanol and dried with N_2_ gas. Each slide was coated with poly-L-lysine, which was applied as a 0.01% w/v solution (Millipore-Sigma) for 20 seconds at room temperature before being extensively rinsed with ultrapure water and dried. Samples were diluted to 20 nM (free DNA and canonical human nucleosomes) or 100 nM (H3-a- and H3-b-containing nucleosomes) using 0.1 µm-sterile filtered dialysis buffer obtained from the final step of nucleosome reconstitution (‘RB2.5’): 10 mM Tris, 2.5 mM NaCl, 5 mM TCEP, 0.2 mM EDTA, pH 7.5. Mass standard calibration was performed using β-amylase diluted to an appropriate working concentration in the same buffer. 60-second videos were recorded in regular image size format (128×34 binned pixels, 18 mm^2^ detection area) and analyzed in Refeyn DiscoverMP (v. 2.3.0). Ratiometric contrast histograms were fitted with Gaussian distributions to extract masses based on the calibration data obtained from the β-amylase standard. Data plots were generated in DiscoverMP and further edited in Adobe Illustrator.

### Thermal Stability Assays

Freshly prepared nucleosomes (containing H3-a, H3-b or *H. sapiens* H3.2) were diluted to 800 nM in solutions containing 10 mM Tris, 2.5 mM NaCl, 5 mM TCEP, 0.25 mM EDTA, pH 7.5, and SYPRO Orange (Invitrogen), diluted to a nominal concentration of 8x from a 5000x stock solution in DMSO. Replicates were transferred into the wells of a 96-well sealable PCR plate and placed in a StepOnePlus Real Time PCR machine (Applied Biosystems). Samples were equilibrated at 25 °C for 1 minute before being subjected to a temperature gradient from 25 to 95 °C, ramping at 1 °C min^−1^. Fluorescence measurements were taken at 1 °C increments using the Cal Orange filter (fluorescence emission maximum, 560 nm). Melting temperatures were estimated from the minima in derivative plots of -d(Fluorescence) / dT versus temperature measured for each sample. All thermal stability data was performed in triplicates. Data analysis and plotting were performed in GraphPad Prism v.10.

### Negative Stain Transmission Electron Microscopy

Nucleosome samples were diluted to 100 nM in RB2.5 (10 mM Tris, 2.5 mM NaCl, 5 mM TCEP, 0.25 mM EDTA, pH 7.5) prior to application on carbon film-coated 400 mesh Cu grids (Electron Microscopy Sciences) glow discharged in a Tergeo-EM plasma cleaner at 15W, 60 seconds. Samples (4 µL) were allowed to incubate on the grid surface for 30s before excess liquid was blotted off with filter paper (Cytiva). Grids were then submerged carbon/sample-side down in 35 µL droplets of 2.5% w/v uranyl formate (Electron Microscopy Sciences) for 5 seconds, blotted, then rinsed 4 times in four additional droplets of uranyl formate solution, before being blotted one last time and allowed to dry in air for 1 minute. Images of negative stain samples were collected at a nominal magnification of 120000x using a FEI Tecnai T12 Spirit transmission electron microscope equipped with a 120 kV LaBr filament (maintained by the BEMS facility at the University of Colorado Boulder). Final images were rescaled and cropped in FIJI^96^.

Reconstruction of 3D densities from negative stain images was performed in CryoSPARC v.4.7.1 (Structura Biotechnology). Briefly, ∼600 micrographs per sample were imported and pre-processed with constant contrast transfer function (CTF) and negative stain settings applied. Particle picking was performed with a blob picker using min/max particle diameters of 100/200 Å, respectively. Particle extraction was performed using a 256’256 px box size (raw pixel size: 2.6 Å px^−1^) and subsequent 2D classification was performed iteratively over three rounds. Final volumes were generated via a single-class *ab initio* reconstruction using 59800 (H3-a) or 28000 (H3-b) particles. Surface representation and editing of the density maps were performed in UCSF ChimeraX v.1.10.1^97^.

### Gradient Fixation, GraFix^98^

Solutions (24 mL) for gradient fixation were prepared as follows: buffer A (20 mM HEPES, 50 mM NaCl, 4 mM TCEP, 0.1 mM EDTA, 10% v/v glycerol, pH 7.5), buffer B (20 mM HEPES, 50 mM NaCl, 4 mM TCEP, 0.1 mM EDTA, 30% v/v glycerol, pH 7.5). Both buffers were filtered through a 0.22 µm pore-size membrane and degassed under vacuum for 30 minutes. Glutaraldehyde (0.15% v/v ∼15 mM; Electron Microscopy Sciences) was added to buffer B just prior to use. To construct the gradient, 7 mL of buffer A was transferred into a 13.2 mL Open-Top Thinwall Ultra-Clear tube (Beckman) using a wide-gauge needle, followed by 7 mL buffer B, displacing the less dense buffer A upwards in the process. Solutions were mixed on a BioComp Gradient Master at a constant angle of 81.5° for 1 minute 54 seconds prior to equilibrating at 4 °C for 30min. Meanwhile, nucleosome samples (containing H3-a or H3-b) were diluted to 3 µM in 200µL RB2.5. In each tube, 250 µL of the top layer of the gradient was removed and replaced with 200 uL of each nucleosome sample. Tubes were transferred into the buckets of a SW40Ti swinging bucket rotor (Beckman Coulter) and subjected to centrifugation at 30000 rpm for 18 hours at 4°C in a Beckman Optima XE90 ultracentrifuge. Samples were then fractionated on the Gradient Master using absorbance readings at 260 nm and 280 nm to track the elution of species suspended in the gradient. Fractions were analyzed by native PAGE, pooled, and dialyzed against 20 mM HEPES, 50 mM NaCl, 5 mM TCEP, 0.2 mM EDTA, pH 7.5 overnight at 4 °C. Final concentrations of crosslinked nucleosomes were measured by DNA absorbance at 260 nm as described above.

### Sample preparation for single particle cryogenic electron microscopy

Glutaraldehyde-crosslinked H3-b-containing nucleosomes (see above) were prepared at a final concentration of 1.27 µM. 4µL samples were mixed with 1µL recombinant AavLEA1 (90 µM; The Histone Source)^99^ right before plunge freezing, rendering final concentrations of ∼1 µM nucleosome and 18 µM AavLEA1. 4 µL of the nucleosome-AavLEA1 mixture were applied to C-Flat 1.2/1.3 300 mesh Cu grids that had been glow discharged in a Tergeo-EM Plasma Cleaner (40 mA, 30 seconds). Plunge freezing was performed in liquid ethane on a Vitrobot Mark IV instrument (Thermo Scientific) with relative humidity, temperature, blot time and relative force set to 100%, 4 °C, 2 and 3 seconds, respectively. Grid quality was first inspected on a 200 kV Glacios 2 TEM (Thermo Scientific) equipped with a Gatan K3 DED camera; successful candidate grids were imaged further on a 300 kV Titan Krios G3i TEM (Thermo Scientific) equipped with a Falcon 4i DED camera. For large scale data acquisition, 10603 exposures were collected at a nominal magnification of 165000x with a raw pixel size of 0.735Å px^−1^, a total exposure dose of 50 electrons Å^−2^, and an applied defocus ranging from −1.8 µm to −0.6 µm (0.2 µm increments).

All cryoEM data processing was performed in CryoSPARC v. 4.7.1 (Structura Biotechnology). Imported exposures were pre-processed via patch motion correction and CTF estimation. None were discarded during manual curation of the motion-corrected micrographs. An initial set of particles was generated with blob picker and extracted with a 400×400 px box size (Fourier cropped to 200×200 px). Iterative 2D classification rendered a curated set of discernible nucleosome-like particles, which were then reextracted from the micrographs (400×400 px box size, without Fourier cropping) and used as inputs for *ab initio* reconstruction using 3 classes. Two of these classes were then used as templates for heterogeneous refinement, from which a single class (54438 particles) was selected for successive rounds of homogeneous refinement, non-uniform refinement, local refinement and local CTF refinement. The resolution of the final map (3.44 Å) was estimated using the gold standard Fourier shell correlation tool.

### Model Building and Refinement

An initial model for the H3-b containing nucleosome was generated from an AlphaFold3 prediction, with inputs being the primary sequences of *H. sapiens* H2A, H2B, H4 and *H. miamia* H3-b, as well as the complementary sequences of each strand of 147DNA. This predictive model was rigidly docked into the unsharpened refined map obtained from the analysis workflow described in the previous section. Real space refinement of the model was performed in PHENIX v.1.21.2^100^ using secondary structural restraints. Multiple iterations of refinement were paired with manual inspection and adjustments to the model made in Coot. Final validation of structural features (clashes, Ramachandran outliers, etc.) and map-model cross-correlation was performed in PHENIX (Supplementary File 5). All figures including the density map and model were prepared in ChimeraX v.1.10.1^97^.

### RNA interference (RNAi)

Double-stranded RNA (dsRNA) was synthesized following previously described protocols^62^. Animals were injected with dsRNA using a Drummond Nanoject II into the gut of the animal. Injections were performed once per day for 3 consecutive days. Animals were amputated transversely at least 3 hours after the final injection and allowed to regenerate. A second round of injections was performed starting on 5 days post-amputation (corresponding to 5, 6, and 7 dpa) and amputated. Control animals were injected with dsRNA targeting *C. elegans unc-22*, absent in *Hofstenia miamia*. Animals were actively monitored for phenotypes or physical defects, such as blastema formation, during the RNAi experiments. Animals were either fixed or harvested for the preparation of downstream processing.

### Bulk ATAC-sequencing library preparation and analysis

ATAC-seq was performed as previously described^101,102^ with the following minor modifications. Blastema regions were isolated from *Hofstenia miamia* tail fragments, placed directly into ice-cold lysis buffer, and homogenized by pipetting. The resulting homogenate was filtered through a 40 µm cell strainer to obtain a single-cell suspension and remove debris, followed by standard ATAC-seq library preparation and sequencing. Biological replicates were performed in triplicate. Downstream processing and analysis, including Genrich and DiffBind^103^, were performed as previously described^101^ with the following modification: all analyses were performed based on a d*e novo* transposable element (TE) library constructed for the PacBio genome assembly using RepeatModeler v.2.0.7^104^, following database preparation (BuildDatabase). Family discovery included the long terminal repeat (LTR) structural discovery module (-LTRStruct), which incorporates LTR_retriever^105^ to more readily identify LTR retrotransposons. Further classification of consensus sequences was supplemented using TEsorter v.1.5.1^106^ against metazoan reference databases, using conserved protein domain homology (hmmscan) with default pass-2 reclassification of unresolved sequence by sequence similarity. The classified repeat library was used to annotate repeat copies genome-wide with RepeatMasker v.4.1.9 using the *de novo* library. To generate a consensus peak set, peaks from all replicates were combined and merged using BEDTools^107^ to produce a unified set of non-overlapping regions. Tornado (heatmap) plots and average accessibility profile plots were generated using deepTools^108^; a signal matrix computed using computeMatrix was plotted using plotHeatmap to display heatmaps, and plotProfile was used to generate the average accessibility profiles. Fragment length distributions were extracted from BAM files using the deepTools bamPEFragmentSize function and visualized with ggplot2 v.3.

### Phylum-level survey of histone H3 sequences

Human canonical histone H3 (NP_003520.1) was used as a query against non-redundant databases to identify histone H3 sequences across selected animal phyla: Annelida, Arthropoda, Cephalochordata, Cnidaria, Ctenophora, Echinodermata, Hemichordata, Mollusca, Nematoda, Placozoa, Platyhelminthes, Porifera, Urochordata, Vertebrata, and Xenacoelomorpha (taxids: 6340, 6656, 7735, 6073, 10197, 7586, 10219, 6447, 6231, 10226, 6157, 6040, 7712, 7742, and 1312402, respectively). Retrieved sequences were filtered to retain hits with an E-value < 1×10^−5^, overall percent identity > 25%, an alignment length of at least 50 amino acids, and a total sequence length between 100 and 150 amino acids. Sequence redundancy was reduced by filtering sequences with 95% identity using CD-HIT^109^. Protein domain and functional annotations were performed using HMMER v.3.3.2 *hmmscan* against the Pfam-A HMM database^110,111^ and InterProScan v.5.77-108.0 against its integrated protein signature databases^49^, respectively. The protein domain annotations were used to define the start and end coordinates for the N-terminal tail (NTT) and histone fold domain (HFD) of each sequence. Percent identities of the annotated NTT and HFD regions were calculated relative to human canonical H3 (NP_003520.1). The 10^th^ percentile of the NTT and HFD percent identity distributions was used as a threshold to partition the H3 sequence dataset into four quadrants. Sequence logos were generated to examine sequence conservation using Jalview v.2.11.5.2^112^. Counts of sequences per animal phylum and divergence class (four quadrants) were compared to expectations under independence using a chi-square test with Monte Carlo-simulated p-values (10^5^ replicates). Standardized Pearson residuals were used to identify over- and under-represented combinations of phylum-class. A t-distributed stochastic neighbor embedding (t-SNE) analysis of histone family sequences was performed using the HFD of H2A, H2B, H3, and H4 family members (n=50 per family), including H3 sequences from the lower-left class. HFD boundaries were defined relative to canonical histone proteins: H3.1 (P68431, residues 65-132); H4 (P62805, residues 32-94); H2A (P04908, residues 28-90); H2B (P62807, residues 39-103; UniProt numbering). HFD sequences were aligned pairwise using BLOSUM62 (gap opening 10, gap extension 0.5) in Biostrings v.2.80.2, and percent identity was calculated relative to the length of the shorter sequence in the pair. The resulting distance matrix (100 minus percent identity) was embedded in two dimensions with Rtsne v.0.17 (theta = 0, perplexity = 10, 2000 iterations, seed #28) and visualized with ggplot2 v.3. Claude Sonnet 5 was used to streamline data processing and visualization.

### Structure prediction and consensus analysis of H3 histone fold domains

HFD sequences from each of the four divergence classes were modeled as monomers using ColabFold v.1.5.5^113^, which runs AlphaFold2^82^ on multiple sequence alignments generated by the MMseqs2 server^114^; no templates were used. Models generated by AlphaFold2 were ranked by mean predicted local distance difference test (pLDDT) score, where the top-ranked model of each sequence was used for subsequent analyses. For each divergence class, models were loaded into PyMOL and superimposed onto the model with the highest mean pLDDT score using the cealign function (combinatorial extension algorithm). To assess structural similarity, all-vs-all C-alpha root mean square deviations (RMSD) were computed between every pair of models using the super function (sequence-independent, structure-based alignment). The consensus structure of each class was defined as a medoid (the model with the lowest RMSD score in each class). The four consensus structures were compared pairwise using US-align^115^. For visualization, consensus structures were superimposed in UCSF ChimeraX v.1.10.1^97^.

## ACKNOWLEDGMENTS

We want to thank the Harvard Medical School Taplin Biological Mass Spectrometry Facility for performing mass spectrometry, the Harvard Bauer core for their assistance in sequencing the ATAC-seq libraries, and the Colorado State University Histone Source core for purifying histone proteins. Thank you to members of the Srivastava and Simoes-Costa labs for thoughtful discussions, including A. Kane, A.P. Kann, J.O. Kimura, C. Breen, O. Duan, R. Busby, and P. Coronel-Córdova for critical reading of the manuscript. Special thanks to D.M. Bolaños for extensive help with the troubleshooting of functional experiments, and to C. Breen for generating heatmaps and volcano plots of published RNA-seq data. We thank the Shared Instrument Pool (RRID: SCR_018986, managed by A. Erbse) at CU Boulder for providing access to ultracentrifuges and mass photometry instruments used in this study (funded by NIH shared instrument grants), and E.W. Hartwick at the BioKEM facility at CU Boulder for assistance with and maintenance of equipment acquiring the cryoEM data reported in this study. We also thank J.M. Allen, Y. Jacob, and L. Banaszynski for scientific insight during data interpretation.

## AUTHOR CONTRIBUTIONS

C.R.L. and M.S. conceived and designed the study. C.R.L, M.L.N., and M.S. wrote the manuscript. C.R.L., M.L.N., S.G.J., P.B., E.M.D., J.D.B., K.L., and M.S. read and edited the manuscript. C.R.L. performed phylogenetic analyses, native histone protein extractions and purifications, FISH, RNAi, confocal imaging, analyses of amplicon-, RNA-, and ATAC-seq data, quantification of heterologous expression images, and survey of histone H3 family sequences across animal phyla. S.G.J. and J.D.B. conducted lentiviral expression and imaging of histone-GFP fusions during heterologous expression experiments. M.L.N and K.L. performed in vitro nucleosome assembly assays, TEM imaging, stability assay, and cryoEM of H3-b nucleosomes. P.B. performed tagmentation and prepared bulk ATAC-seq libraries. E.M.D., and K.L., and M.S. provided scientific insight during data analysis.

---

**Supplementary Figure 1.**
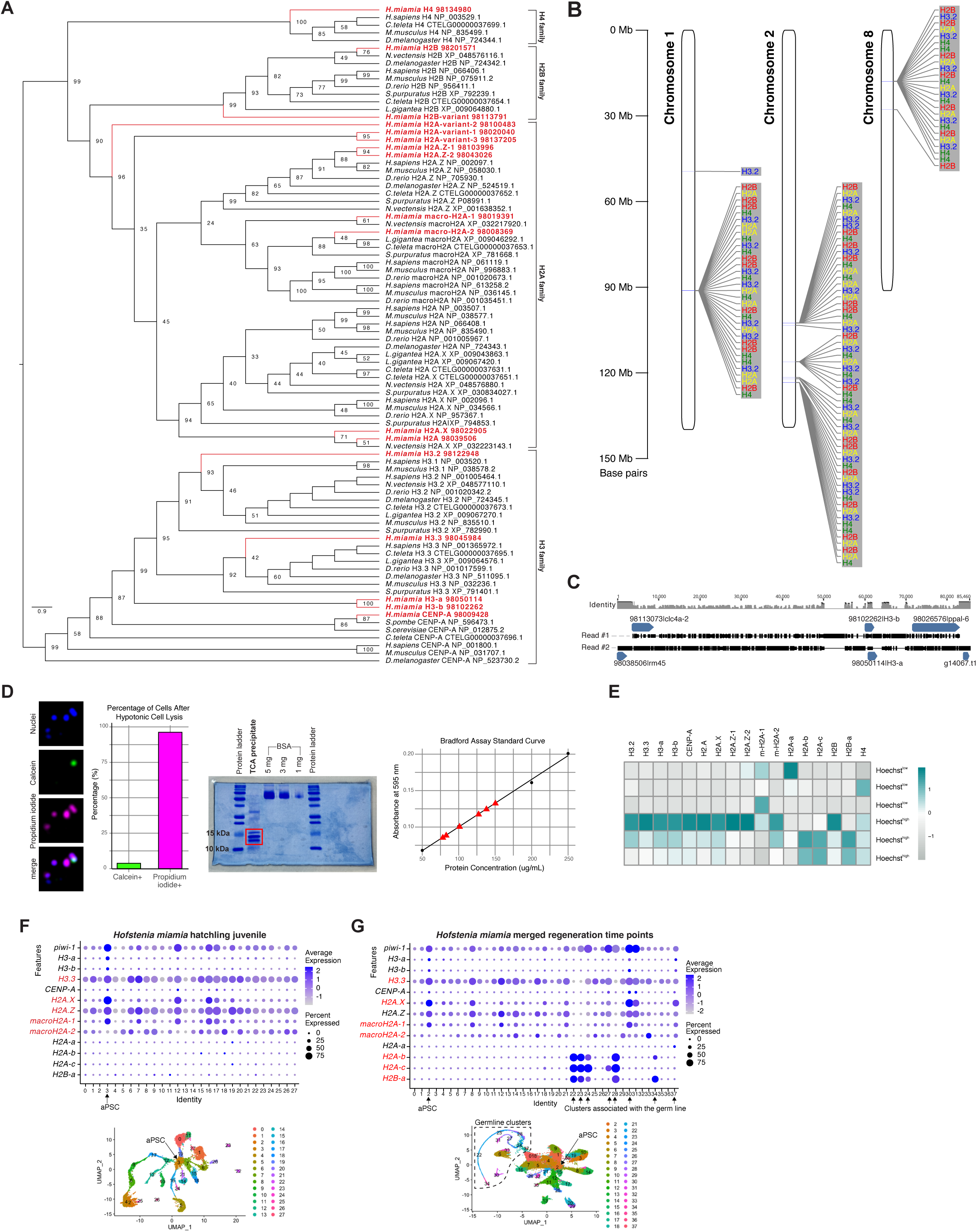
Histone family phylogenies, genomic organization, and nuclear histone purification in *H. miamia*. (A) Maximum-likelihood phylogenetic tree for homologs of the H2A, H2B, H3 and H4 families with bootstrap support shown at each node. The *H. miamia* histone homologs are highlighted in red and fall in highly supported clades with known orthologs of each respective histone family. (B) Chromosome maps showing canonical histone gene clusters on chromosomes 1, 2, and 8. Individual histone genes are color-coded: H3.2, blue; H4, green; H2A, yellow; H2B, red. Genomic coordinates are provided in Supplementary File 1. (C) Alignment of two independent PacBio long reads spanning H3-a, H3-b, and their respective flanking regions confirmed that H3-a and H3-b are encoded at distinct genomic loci. Pairwise identity is indicated above the alignment. (D) Purification of histone proteins from isolated *H. miamia* nuclei. *Left*: Hypotonic lysis ruptured ∼95% of cell membranes, enabling nuclei isolation. Calcein labels intact cells (green) and propidium iodide labels lysed cells (magenta). *Center*: Trichloroacetic acid (TCA) precipitation enriched nuclear proteins in the expected histone size range (10-15 kDa, red box), as assessed by sodium dodecyl sulfate-polyacrylamide gel electrophoresis; indicated gel region was excised for downstream mass spectrometry. Bovine serum albumin (BSA) was included as reference. *Right*: Bradford assay measurements yielded protein concentrations of 80-150 µg/mL (red triangles), sufficient for mass spectrometry. (E) Heatmap showing relative expression of *H. miamia* histone genes in Hoechst^high^ vs Hoechst^low^ cell populations^44^. H2A.Z-1 and H2A-a had insufficient counts for q-value estimation in the likelihood ratio test and were therefore excluded from the volcano plot in the main figure (see Supplementary File 3). (F-G) Dot plots showing histone variant expression across *H. miamia* hatchling juvenile cell populations (F) and merged regeneration time points (G) from Hulett et al., 2023. Dot size represents the proportion of cells expressing each histone variant, and dot color indicates average expression. Histone variants highlighted in red are expressed in cell clusters outside the aPSC population. UMAPs below each dot plot show the corresponding cell clusters; the dashed box indicates germline clusters.

**Supplementary Figure 2.**
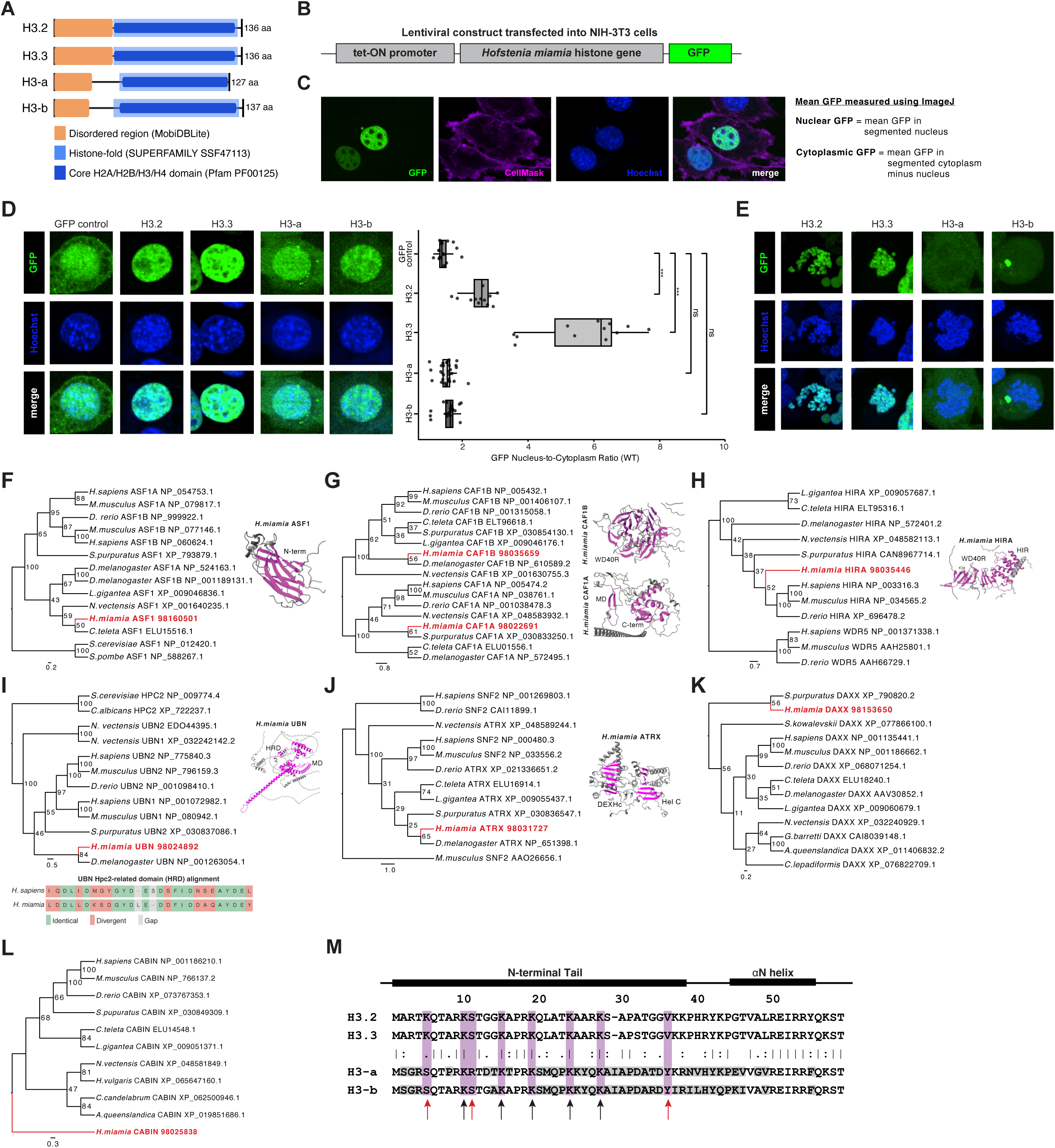
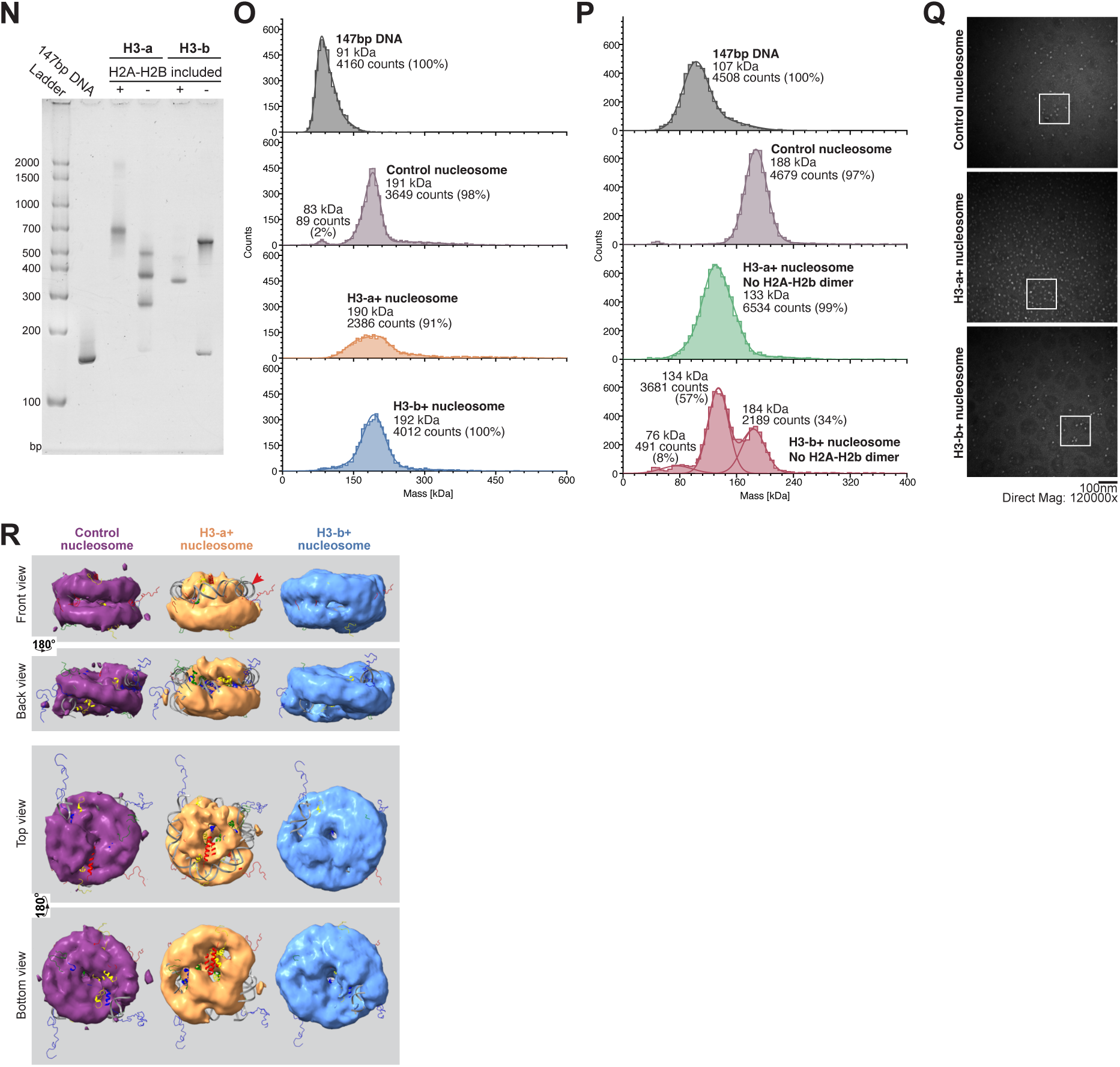
Sequence features, chaperone recognition, and nucleosome assembly of H3-a and H3-b. (A) Predicted domain organization of the *H. miamia* H3 family based on InterProScan, showing disordered regions (orange) and HFDs (light blue and dark blue). (B) Schematic of the lentiviral construct used for heterologous expression of GFP-tagged *H. miamia* H3 proteins under a Tet-ON promoter. (C) Representative segmentation of NIH-3T3 used to quantify nuclear and cytoplasmic GFP signals. CellMask and Hoechst were used to define cell and nuclear boundaries, respectively; quantification was performed in Fiji^96^. (D) Heterologous expression of GFP-tagged *H. miamia* H3 proteins in NIH-3T3 cells. Representative confocal images are shown (left), with quantification of nucleus-to-cytoplasm GFP signal relative to GFP control (right). H3.2 and H3.3 show strong nuclear localization, whereas H3-a and H3-b show reduced nuclear enrichment. ***, *p*-value < 0.001; ns, not significant. (E) Localization of GFP-tagged *H. miamia* H3 proteins in colcemid-arrested NIH-3T3 cells. H3.2 and H3.3 co-localize with condensed chromatin, whereas H3-a and H3-b show reduced chromatin association. (F-L) Maximum likelihood phylogenetic analyses of *H. miamia* homologs of histone chaperone components: ASF1 (F); CAF1A and CAF1B (G); HIRA (H); UBN (I); ATRX (J); DAXX (K); and CABIN (L). Predicted structures of representative conserved domains are shown alongside each tree. Homologs of ASF1, CAF1A/CAF1B, HIRA, UBN, and ATRX were recovered within supported orthologous clades, whereas candidate DAXX and CABIN sequences showed less confident phylogenetic placement. The predicted Hpc2-related domain of *H. miamia* UBN shares 57.7% identity with its human counterpart (I). No clear HJURP homolog was identified. Domain abbreviations: WD40R, WD40 repeat; HIR, HIRA domain; HRD, Hpc2-related domain; MD, middle domain; DEXHc, DEAH-box helicase C-terminal domain; Hel C, helicase superfamily C-terminal domain. (M) Multiple sequence alignment of respective *H. miamia* H3 NTTs and ɑN helix. Residues corresponding to known post-translational modification sites in canonical H3 are highlighted in magenta. Conserved sites are indicated by black arrows, substitutions in H3-a or H3-b by red arrows, and divergent flanking residues by gray shading. H3-a and H3-b retain several canonical lysines, including K9, K14, K18, K23 and K27, while substitutions occur at other conserved positions, including K4 and K36. (N) Native polyacrylamide gel electrophoresis analysis of complexes assembled with H3-a or H3-b in the presence or absence of H2A-H2B dimers. Complete histone complements produce homogeneous nucleosome-sized complexes, whereas omission of H2A-H2B results in heterogenous assemblies and residual free DNA. DNA size markers are indicated in base pairs (bp). (O-P) Mass photometry of particles assembled in the absence (O) or presence (P) of H2A-H2B dimers. Particle populations are resolved by molecular mass (kDa), with relative abundances indicated. (Q) Representative negative-stain TEM micrographs of reconstituted nucleosomes. White boxes indicate the insets presented in Figure 2E. Scale bar, 100 nm; magnification, 150,000x. (R) Front, Back, Top and Bottom views of low-resolution density maps reconstructed from H3-a- and H3-b-containing nucleosomes, overlaid with the canonical nucleosome structure (PDB: 1KX5). The red arrow indicates unresolved density in the H3-a+ nucleosome.

**Supplementary Figure 3.**
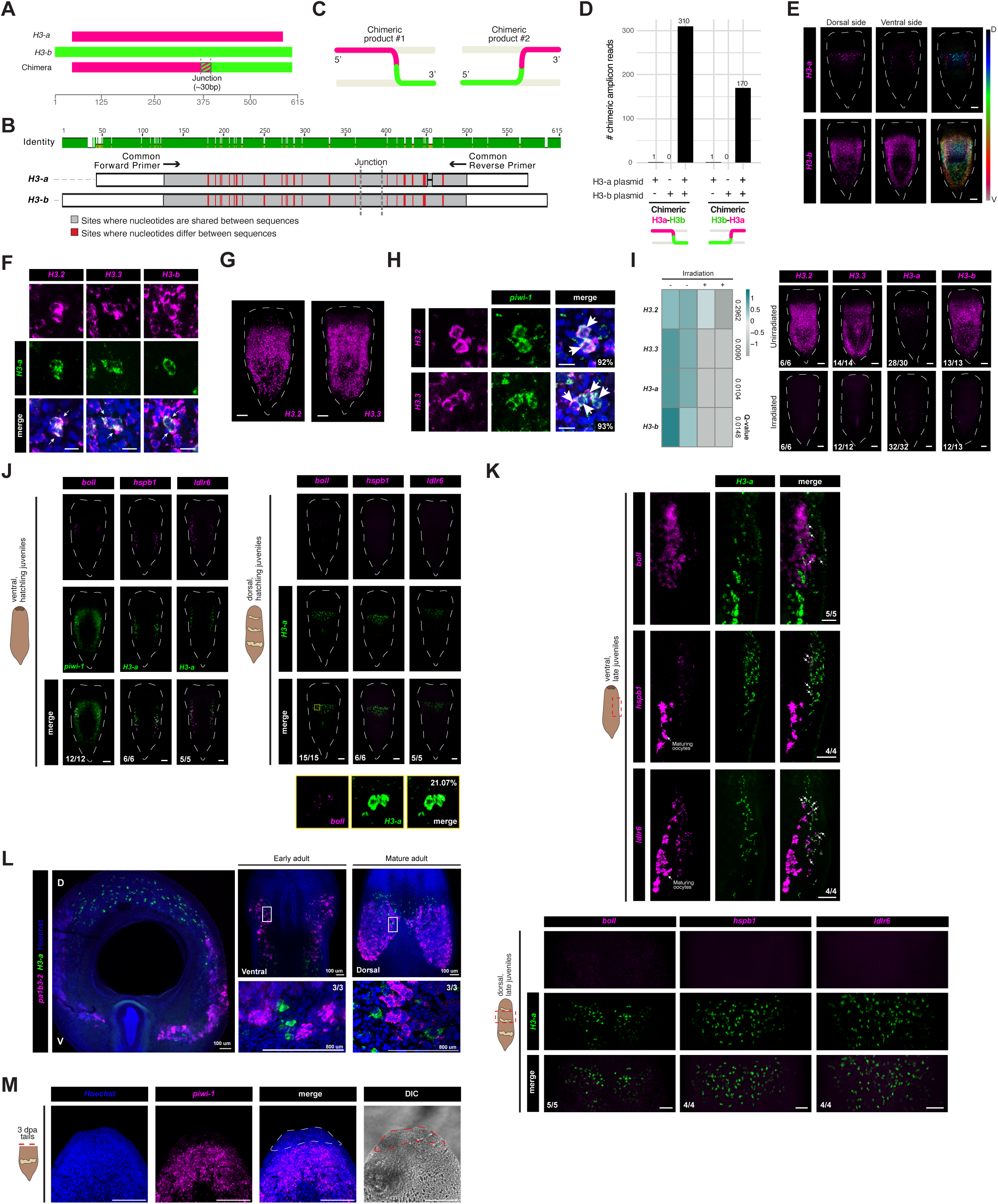
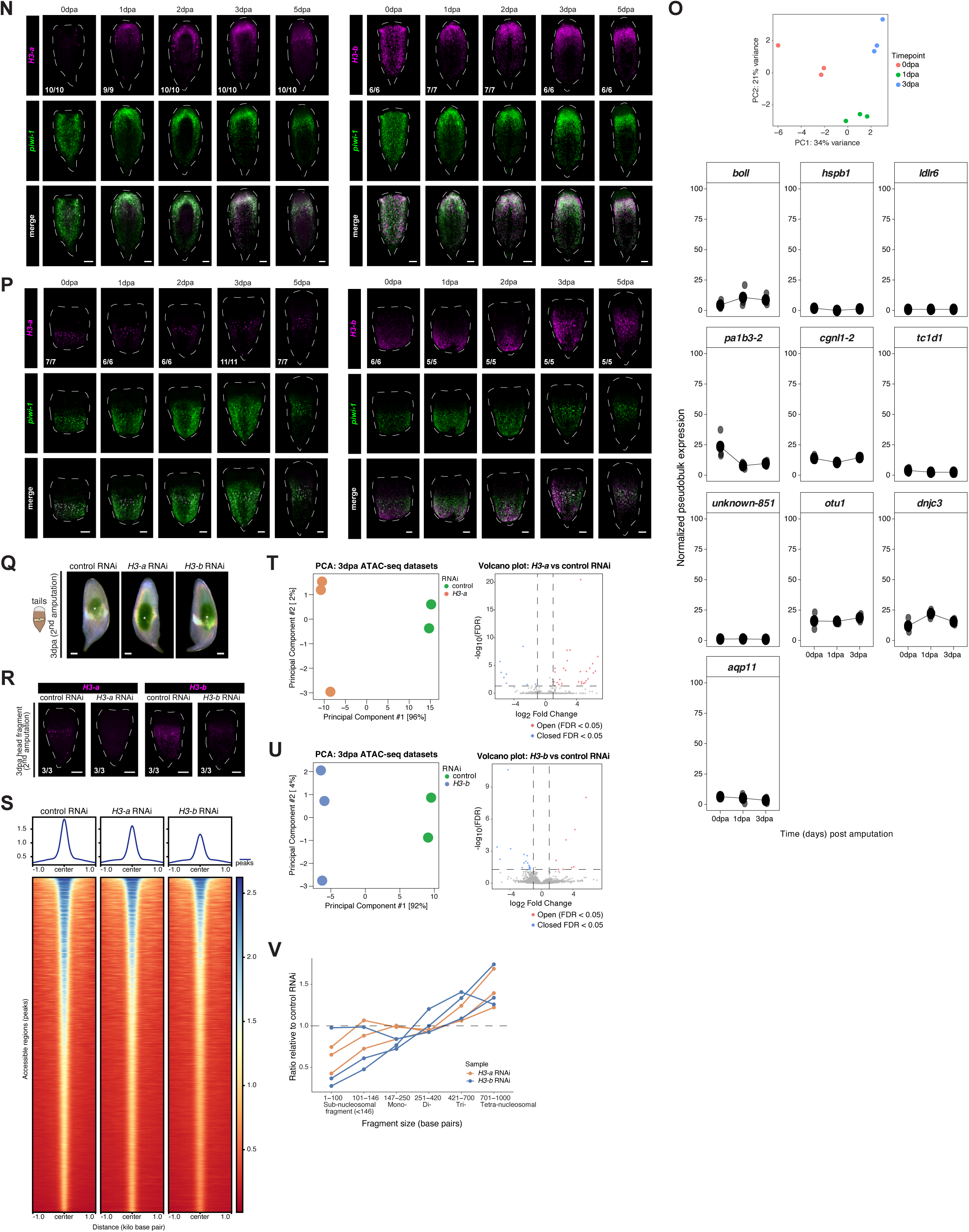
Validation and extended characterization of H3 expression and chromatin accessibility in *H. miamia* aPSCs. (A) Schematic alignment of the *H3-a* and *H3-b* sequences from this study with the *H3-a* clone from our previous study^17^, revealing a chimeric sequence in which the 5’ region of *H3-a* is fused to the 3′ region of *H3-b* (denoted here as the “junction” region; see Supplementary Figure 2). Scale indicates nucleotide position. (B) PCR amplification of *H3-a* and *H3-b* individually or as a 1:1 mixture using shared primers to test for chimera formation during amplification^117–121^. Single-template reactions serving as controls. (C) Schematic of possible reciprocal H3-a/H3-b and H3-b/H3-a chimeras resulting from template switching at the junction region. (D) Amplicon sequencing identified chimeric products in mixed-template reactions, supporting a PCR-derived origin of the previously cloned sequence. A newly designed, sequence-verified *H3-a*-specific probe revealed both the previously reported dorsal *H3-a*+ subpopulation^17^ and an additional ventral *H3-a*+ subpopulation. This new probe was used for all imaging and functional experiments in this study. (E) Maximum-projection FISH images showing the dorso-ventral distributions of *H3-a* and *H3-b* in intact animals. Z-stacks were divided to visualize dorsal and ventral cell populations separately. Scale bars, 100 µm. (F) Double FISH for *H3-a* with *H3.2*, *H3.3*, and *H3-b* in intact animals. Co-expressing cells are indicated by white arrows. Scale bars, 10 µm. (G) FISH for *H3.2* and *H3.3* in hatchling juveniles. Scale bars, 100 µm. (H) Double FISH for *H3.2* and *H3.3* with *piwi-1* in intact animals. Co-expressing cells are quantified and indicated by white arrows. Scale bars, 10 µm. (I) Irradiation sensitivity of H3 family gene expression in animals exposed to 10,000 rads. *Left*: Relative H3 family gene expression in irradiated and control animals measured by bulk RNA-seq; q-values from likelihood ratio tests are indicated^44^. *Right*: Representative FISH for *H3.2*, *H3.3*, *H3-a*, and *H3-b* 7 days post-irradiation. Scale bars, 100 µm; numbers indicate sample size. (J) Double FISH for *H3-a* with *boll*, *hspb1*, or *ldlr6* in ventral or dorsal regions of hatchling juveniles. Ventral *H3-a*+ cells co-express all three markers, whereas *hspb1* and *ldlr6* are not detectably expressed dorsally. Double FISH for *boll* and *piwi-1* confirms their co-expression. *Inset*: Faint, low-level dorsal *boll* expression co-localizes with a small subset of *H3-a*+ cells (21.07% of *H3-a*+ cells were *boll*+, n=3). Scale bars, 100 µm; numbers indicate sample size. (K) Double FISH for *H3-a* with *boll*, *hspb1*, or *ldlr6* in ventral (*top*) or dorsal (*bottom*) regions of late juveniles. Ventral *H3-a*+ cells co-express these germline-associated markers, whereas dorsal *H3-a*+ cells do not. Scale bars, 100 µm; numbers indicate sample sizes. (L) Double FISH for *H3-a* and *pa1b3-2* in maturing adults. *H3-a*+ cells are positioned near, but do not detectably co-express *pa1b3-2*. Scale bars are indicated in each panel; numbers indicate sample sizes. (M) Representative FISH and DIC images at 3 dpa blastema showing sub-blastemal *piwi-1* expression. Dotted outline indicates the blastema. Scale bars, 100 µm. (N) Representative double FISH for *H3-a* or *H3-b* with *piwi-1* in regenerating tail fragments at 0-5 dpa. Samples correspond to those shown in Figure 3G. Scale bars, 100 µm; numbers indicate sample sizes. (O) Normalized pseudobulk expression of germline-associated genes in the aPSC cluster at 0, 1, and 3 dpa using regeneration scRNA-seq data from Hulett et al., 2023. Principal component analysis (PCA) shows that samples cluster by regeneration time point. (P) Representative double FISH for *H3-a* (*left*) or *H3-b* (*right*) with *piwi-1* in regenerating head fragments at 0-5 dpa. Scale bars, 100 µm; numbers indicate sample sizes. (Q) Representative regenerating tail fragments following control, *H3-a* and *H3-b* RNAi, imaged 3 days after the second amputation. White asterisks indicate residual injection dye. Scale bars, 100 µm. (R) Representative FISH confirming depletion of *H3-a* or *H3-b* transcripts following the corresponding RNAi treatment. Scale bars, 100 µm; numbers indicate sample sizes. (S) ATAC-seq signal centered on consensus peak midpoints (±1 kb) across control, *H3-a* and *H3-b* RNAi conditions. Heatmaps show normalized read density for individual accessible regions ordered by control signal; aggregate profiles above show mean accessibility across all regions. (T-U) Principal component and differential accessibility analysis following *H3-a* (T) or *H3-b* (U) RNAi. *Left*: PCA of normalized ATAC-seq read counts across consensus peaks; each point represents a biological replicate. *Right*: Volcano plots of differential analysis relative to control RNAi. Each point represents an accessible region plotted by log_2_ fold change and -log_10_(FDR); significantly differentially accessible regions (FDR < 0.05) are highlighted. (V) Per-sample ATAC-seq fragment-size distributions following control, *H3-a*, or *H3-b* RNAi. Ratios relative to the mean of control samples are plotted across fragment lengths.

**Supplementary Figure 4.**
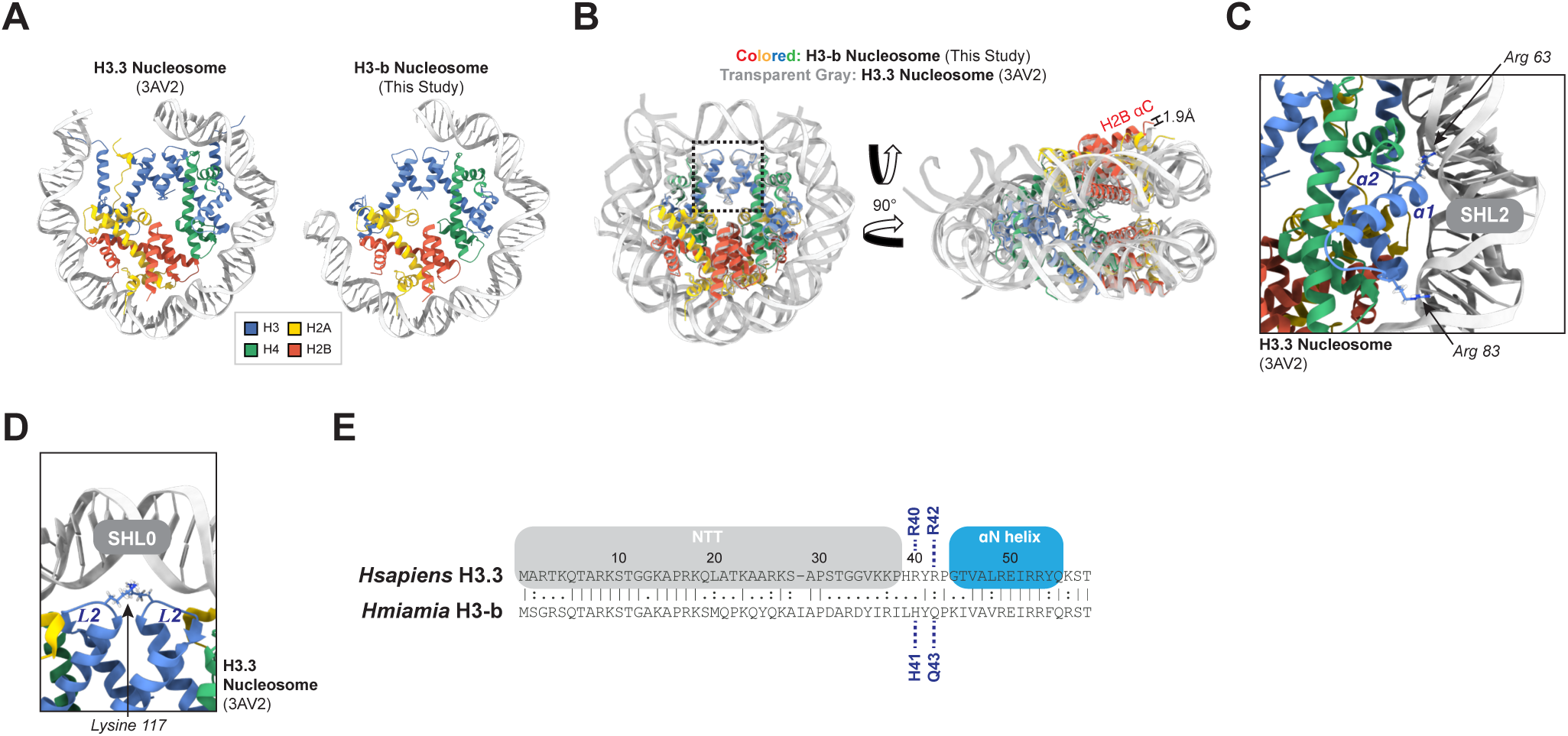
Histone-DNA interactions in the canonical H3.3-containing nucleosome. (A) ‘Half-view’ comparison of the H3.3-(PDB: 3AV2) and H3-b-containing nucleosome structural models. One H2A-H2B dimer, a single H4 monomer, and 72 base pairs (H3.3) or 58 base pairs (H3-b) of DNA are hidden for clarity. (B) Overlay of the H3-b-containing nucleosome (colored ribbon) and H3.3-containing nucleosome (transparent gray) structural models, aligned on the H3-H3’ four-helix bundle (4HB; black hatched box). Distance was measured between the Cɑ of H2B residues Lys86 and Lys120 in the H3-b and H3.3 structures, respectively. (C) Enlarged view of the H3.3 ɑ1-loop1-ɑ2 region in the H3.3-containing nucleosome, highlighting Arg63 and Arg83 (*H. sapiens* numbering) near superhelical location 2 (SHL2) of nucleosomal DNA, corresponding to the view shown in Figure 4C. (D) Enlarged view of the nucleosomal dyad (SHL0), showing the Lys117 (*H. sapiens* numbering) sidechains from both H3.3 monomers and their positions relative to DNA, corresponding to the view shown in Figure 4D. Histone proteins are color-coded: H3.3, blue; H4, green; H2A, yellow; H2B, red. (E) Pairwise alignment of the first 59-60 residues of the *H. miamia* H3-b and *H. sapiens* H3.3 primary sequences. Colored boxes highlight the positions of the N-terminal tail (NTT) and ɑN helix in H3.3.

**Supplementary Figure 5.**
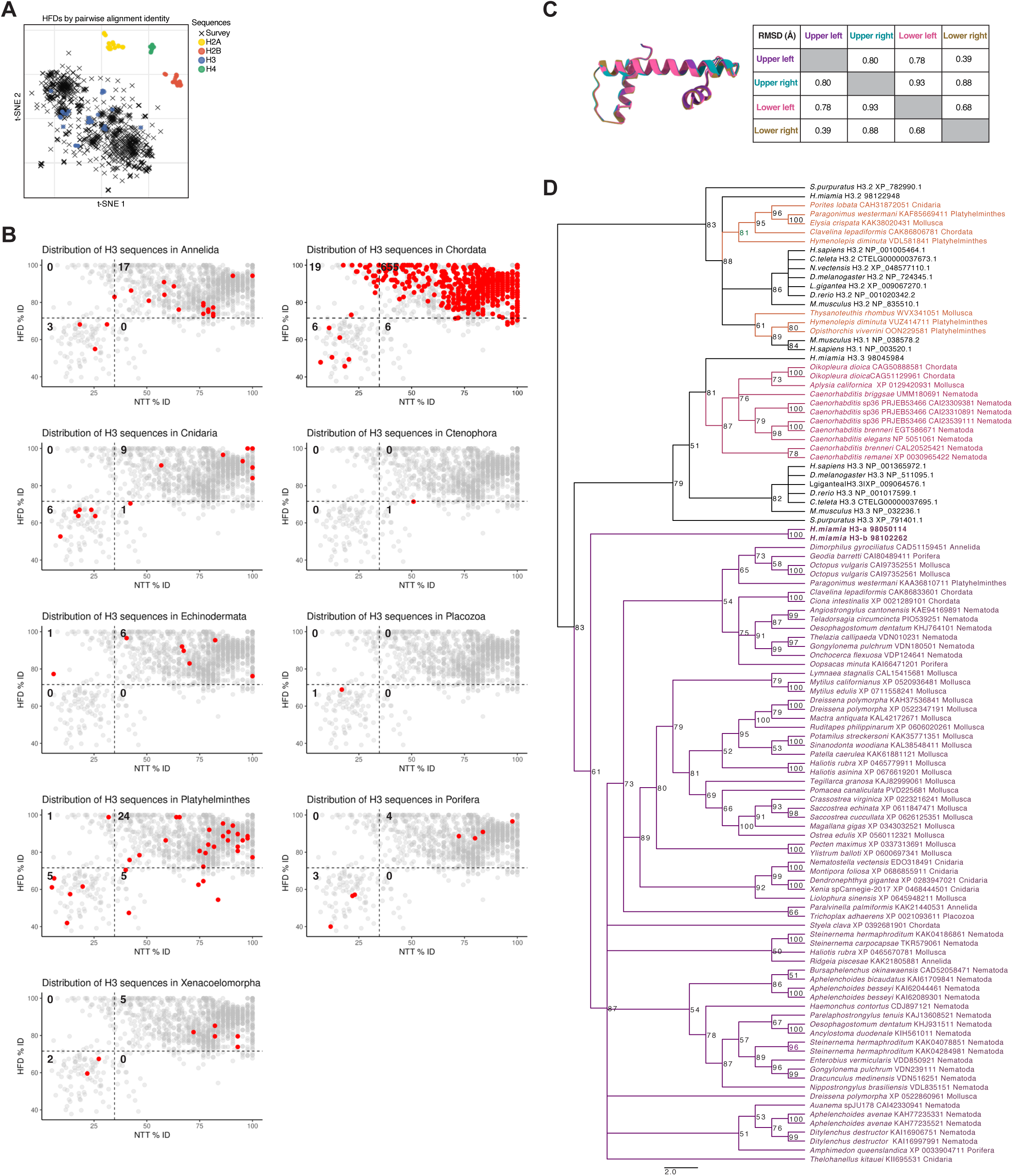
Sequence, structural, and phylogenetic characterization of divergent animal H3 proteins. (A) t-SNE embedding derived from pairwise HFD sequence distances among H2A (yellow), H2B (red), H3 (blue), and H4 (green) proteins. Black crosses represent identified H3 sequences in the cross-phyla survey, which cluster with the H3 family. Pairwise distances were calculated as 100 minus percent sequence identity from BLOSUM62-based alignments. (B) Distribution of annelid, chordate, cnidarian, ctenophore, echinoderm, placozoan, platyhelminth, poriferan, and xenacoelomorph H3 sequences across divergence classes. Sequences from the indicated groups are highlighted in red, with all other H3 sequences shown in gray. Numbers indicate sequence counts within each quadrant. (C) Superimposition of class-representative medoid (consensus) HFD structures predicted by AlphaFold2 for the upper-left (purple), upper-right (blue), lower-left (pink), and lower-right (yellow) classes. The table reports pairwise core Cɑ RMSD (Å) between medoid structures. (D) Maximum-likelihood phylogenetic tree of lower-left class H3 sequences together with representative canonical H3.1/H3.2 and H3.3 proteins. Node values indicate bootstrap support.

**Supplementary File 1. Histone and histone-chaperone annotations in *Hofstenia miamia*.** Protein and nucleotide sequences for the annotated *H. miamia* histone and histone-chaperone complement, genomic coordinates of canonical histone genes, and R-scape analyses used to identify conserved 3’ UTR stem-loop structures in canonical histone transcripts.

**Supplementary File 2. Sequence alignment of H3-a, H3-b, and the previously identified chimeric H3 sequence.** Multiple sequence alignment of *H3-a* (yellow), *H3-b* (blue), and the previously cloned chimeric H3 sequence, which is consistent with a PCR amplification artifact generated from *H. miamia* cDNA. The H3-a/H3-b junction is highlighted in green.

**Supplementary File 3. Differential expression of *H. miamia* histone genes in Hoechst^high^ and Hoechst^low^ cell populations.** Estimated read counts, TPM values, and differential expression statistics for *H. miamia* histone genes in Hoechst^high^ and Hoechst^low^ samples. H2A.Z-1 and H2A-a had insufficient counts for q-value estimation in the likelihood ratio test and are reported as NA.

**Supplementary File 4. Chromatin-accessibility and fragment-size analyses following *H3-a* or *H3-b* RNAi.** Differential ATAC-seq accessibility statistics for control versus *H3-a* or *H3-b* RNAi, including DiffBind output and fragment-size distribution counts.

**Supplementary File 5. Cryo-EM data collection, processing, refinement, and validation statistics for the H3-b-containing nucleosome.** Single-particle cryo-EM collection and processing parameters, map-resolution statistics, atomic model refinement, map-to-model correlations, model composition, geometric deviations, and structural validation metrics for the GraFix-crosslinked H3-b-containing nucleosome structure.

**Supplementary File 6. Cross-phyla survey of animal histone H3 sequence divergence.** H3 sequences recovered across animal phyla, NTT and HFD percent identities, divergence-class assignments, taxonomic annotations, and chi-squared analyses of class distributions, including observed and expected counts and fold enrichment by phylum.

